# A self-amplifying microbial–abiotic sulfur relay drives field feasible bauxite residue remediation

**DOI:** 10.64898/2026.08.20.746091

**Authors:** Jing Zhao, Julian Zaugg, Fang You, Narottam Saha, David Parry, Philip Hugenholtz, Longbin Huang

## Abstract

Bauxite residue (BR), the haloalkaline byproduct of alumina refining, represents the largest and most costly environmental challenge facing the global aluminium industry, yet sustainable remediation has remained elusive because no rapid and field-feasible technology can overcome its recalcitrant alkalinity. Here, we establish a self-amplifying microbial–abiotic sulfur relay that drives rapid *in situ* acid generation and sustained dealkalization of BR across laboratory and glasshouse experiments and a field trial, where dealkalized residue subsequently supported spontaneous pioneer-plant colonization. Mechanistic assays and multi-omics analyses show that the relay is initiated by microbial reduction of elemental sulfur (S_8_) to HS^−^ under oxygen-limited conditions. The resulting HS^−^ abiotically attacks and solubilizes solid S_8_, generating a mobile pool of polysulfides (S□²□). In anoxic microsites, polysulfide reduction regenerates HS^−^, which mobilizes additional S_8_ and amplifies sulfur turnover by increasing sulfur mobilization and bioavailability. In oxic microsites, 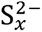 are abiotically converted to thiosulfate and reactive S^0^, which are subsequently microbially oxidized to sulfate and acidity. By coupling biotic reductive initiation and regeneration with abiotic sulfur mobilization and oxidation, followed by biotic terminal oxidation, this relay overcomes the low bioavailability of S_8_ and the constraints of extreme haloalkaline conditions, providing a low-cost, field-feasible strategy for efficient and sustained BR remediation.

## Introduction

Toxic tailings from global mining represent a planetary-scale problem, with hundreds of billions of tonnes accumulated^1^ and production increasing by 70 billion tonnes annually^2^. Among these, bauxite residue (BR), a by-product of alumina refining, is one of the largest and most hazardous mining wastes. It is extremely fine (>90% of particles <10 μm), strongly alkaline (pH >11), sodic (>50 g kg□ ¹), and heavily buffered (∼30 g kg□ ¹ CaCO□ equivalent), a combination that resists neutralization and precludes ecological reuse^3^. Global inventories were estimated at ∼4.6 billion tonnes in 2018^4, 5^ and continue to grow by approximately 175 million tonnes annually^6^, posing long-term risks to water, soil, and public health. The global aluminum industry is facing an urgent challenge to achieve sustainable BR management and remediation^7^.

A key prerequisite for sustainable BR rehabilitation is irreversible dealkalization and neutralization, which remove the constraint of extreme pH and facilitate the mineralogical and geochemical stabilization of metal(loid)s (e.g., Al, As, Cu, Mo, V, Zn), enabling *in situ* soil formation and revegetation^8–11^. The high alkalinity of BR arises from two principal sources: (i) residual caustic compounds (e.g., OH^−^) left behind from the Bayer process, and (ii) strongly buffered alkalinity resulting from alkali ions structurally bound within insoluble aluminosilicate minerals such as sodalite^12, 13^. Conventional neutralization strategies, including capping, seawater flushing^14^, gypsum amendment, mineral acids, and acid gas treatments^3^, have been widely explored, yet remain constrained by cost, scalability, and inconsistent long-term performance^15, 16^. Specifically, seawater neutralization, which is geographically constrained, and gypsum addition primarily transform soluble alkalinity into secondary mineral phases rather than eliminating the underlying alkaline reservoir. Consequently, treated residues often remain alkaline (pH ∼9–10) due to pH rebound^17^. Such conditions can maintain trace elements in soluble, weakly sorbed forms, limiting their long-term stabilization^18^. More aggressive treatments, such as direct addition of acid, can rapidly reduce pH but are costly, difficult to scale, and environmentally risky. These limitations highlight the need for environmentally friendly strategies capable of achieving sustained and irreversible dealkalization.

Continuous, rapid *in situ* acid generation driven by microbial activity offers an alternative, potentially sustainable route for BR dealkalization. However, fermentation of organic matter yields only weak organic acids and is constrained by environmental factors such as carbon availability and moisture, limiting its effectiveness in extremely alkaline conditions^13, 19–22^. Microbial elemental sulfur (S_8_) oxidation has been explored for acidifying alkaline soils (pH ∼8–9)^23^ and iron ore tailings^24^, but under BR’s extreme haloalkalinity (pH >10), hydrophobic S_8_ is poorly bioavailable and microbial activity is physiologically constrained, such that direct S_8_ oxidation has failed or proceeded far too slowly to be useful^25^. Here, we overcome these limitations by establishing a self-amplifying microbial–abiotic sulfur relay in redox-heterogeneous BR that bypasses the slow direct oxidation of S□ and the constraints imposed by extreme haloalkaline conditions, enabling rapid in situ sulfuric acid generation that drives BR dealkalization to the point of spontaneous plant colonization and ecological reuse, as validated across laboratory, glasshouse, and field trials. Mechanistic experiments, together with metagenomic and metatranscriptomic analyses, show that the relay begins with a limited reductive flux, in which S_8_ is microbially reduced to a small HS□ pool that chemically attacks solid S_8_, dissolving it into a mobile, bioavailable pool of active polysulfides. In anoxic microsites, these intermediates are readily reduced back to HS□, mobilizing more S_8_ ; in oxic microsites, they are rapidly and spontaneously oxidized to thiosulfate and newly formed S□, both of which serve as substrates for microbial oxidation to sulfate and acid generation. In field applications, the self-amplifying microbial–abiotic relay is readily activated by co-amendment with S□ and organic matter, which create heterogeneous redox conditions and supply organic substrates for the initial S□ reduction step. Organic matter also contributes organic acids that further accelerate dealkalization and nutrients that promote plant colonization. Overall, our findings establish a mechanistically informed and field-validated pathway for irreversible BR dealkalization, providing a scalable and environmentally sustainable strategy for remediating one of the world’s largest alkaline waste streams.

## Results

### Lab-scale proof-of-concept

Laboratory incubations established that co-amendment with organic matter (OM) and elemental sulfur (S_8_) enhances bauxite residue (BR) dealkalization relative to single OM amendments or unamended controls (**Supplementary Fig. 2a**). L_OM3S (40% v/v mulch + 3% w/w S_8_) produced the largest pH decrease (1.93 ± 0.07 units), reaching pH 8.86 ± 0.07 after 48 days. This decline occurred in two phases: a rapid decrease of 1.10 ± 0.02 units within 24 hours, consistent with release of organic acids from the mulch amendment, followed by a sustained decline (0.018 units day□¹) tracking sulfur transformation to sulfate (**Supplementary Fig. 2b**). The L_OM3S treatment showed the highest rate of oxidized-sulfur (247.5 mg L□¹ day□¹) and sulfate (98.7 mg L□¹ day□¹) production among all treatments, approximately double the rate of oxidative sulfur accumulation in the S□-only treatment (L_3S), indicating that organic matter amendment strongly enhanced sulfur transformation (**Supplementary Fig. 2c**).

### Glasshouse validation of sustained dealkalization

A glasshouse trial (G_, 40% OM with 0–3% S_8_) tested dealkalization performance under more field-relevant conditions (**Fig. 1a**). The co-amended treatment with 3% S_8_ (G_OM3S) produced the greatest and most sustained pH reduction, decreasing by 1.68 ± 0.42 units to pH 8.90 ± 0.30 by day 282 (**Fig. 1b**). By contrast, lower- or no-S_8_ treatments fell only modestly and stabilized near pH 10 within the first week (**Fig. 1b**). At the end of the experiment (Day 282), treated tailings were collected for further analysis. The pH□:□ (pH of a 1:5 tailing-to-water suspension, reflecting the soluble fraction of alkalinity) decreased by 1.07 units in G_OM3S (**Fig. 1c**), while solid-phase alkalinity declined by 0.48 mmol g**□** ¹ relative to untreated BR (**Fig. 1d**). Mineralogical analysis (qXRD) showed an 11.3% reduction in sodalite content (**Supplementary Fig. 3**), indicating the treatment depletes the mineral alkalinity reservoir. G_OM3S also raised nutrient availability with increased total organics (**Fig. 1h**) and nitrogen (**Fig. 1j**) in porewater and total organic carbon in treated tailings (**Fig. 1m**), and reduced sodality with the lowest exchangeable sodium percentage (ESP) (**Fig. 1n**) and sodium adsorption ratio (SAR) (**Fig. 1o**), indicating its potential to transform BR into a technosol suitable for plant growth.

**Figure 1:**
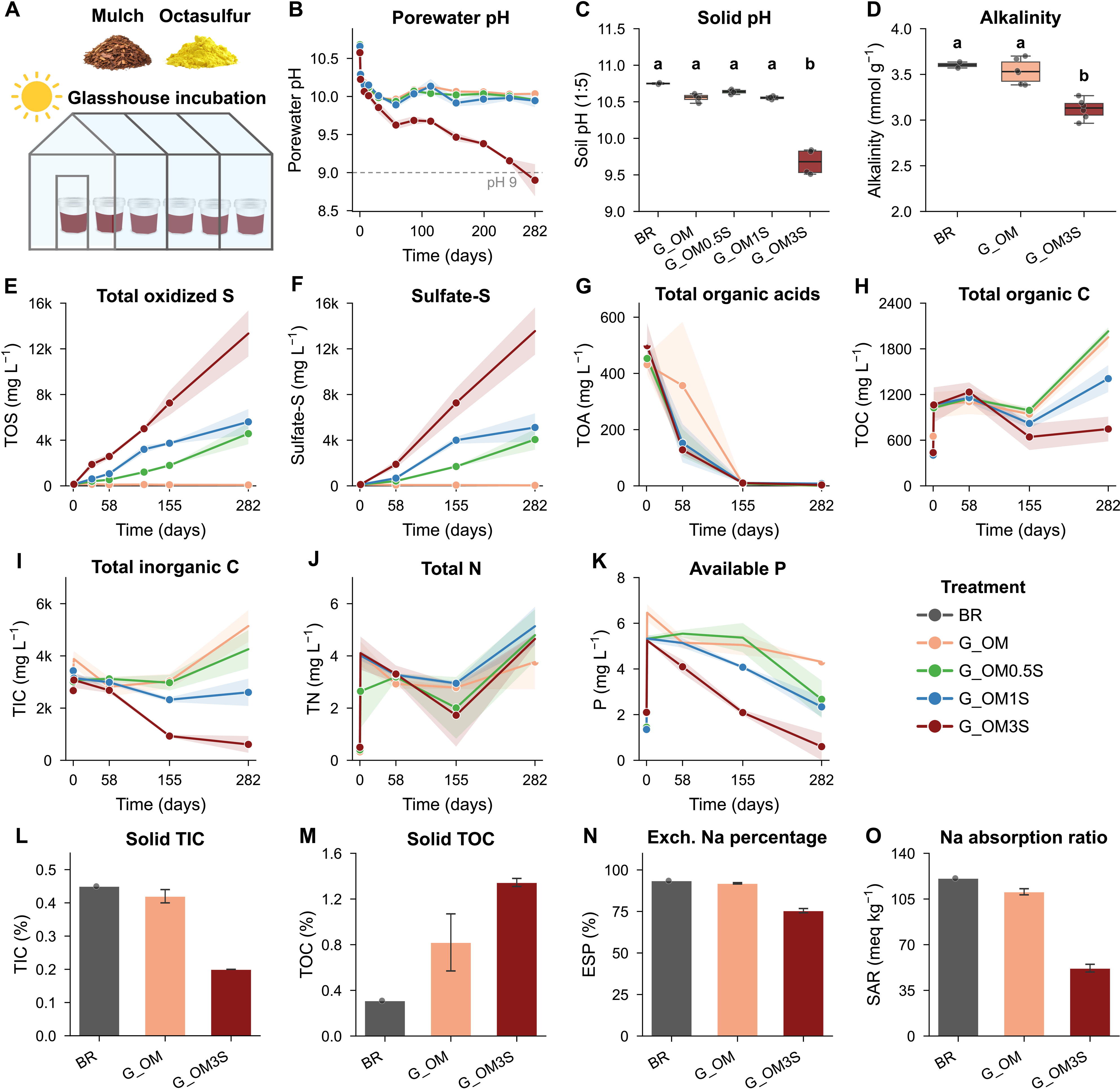
**(a)** Experimental schematic. Bauxite residue was amended with organic matter (OM) alone or with OM plus elemental sulfur (S_8_) at 0.5%, 1%, or 3% (w/w) and incubated in a glasshouse for 282 days (temperature 10-30 degrees C; soil saturation decreasing from 70% to 30%). (**b)** Porewater pH over time; dashed line marks pH 9. **(c, d)** Final-day tailings pH (1:5 solid:water) and alkalinity. **(e–k)** Porewater dynamics of total oxidized sulfur, sulfate, total organic acids, total organic carbon, total inorganic carbon, total N, and available P. **(l–o)** Final-day solid-phase TIC, TOC, exchangeable sodium percentage, and sodium adsorption ratio. Data show means ± s.e. (time series, n = 2 pots per treatment per time point) or box plots/means as indicated.

The temporal pattern of dealkalization in G_OM3S mirrored that observed in laboratory incubations, with an initial rapid dealkalization phase driven by organic acids (**Fig. 1g**), followed by a sustained decline associated with sulfate generation (**Fig. 1f**). High S_8_ loading and the associated increase in sulfate production were also linked to accelerated total organic carbon (TOC) depletion, suggesting a coupling of carbon and sulfur metabolism. Compared to laboratory incubations, the glasshouse system showed greater conversion of sulfur to sulfate. This difference is consistent with improved oxygen availability in the amended BR, facilitated by the addition of coarse mulch and cyclic moisture conditions. Mulch amendment reduced bulk density (from 1.7–1.9 to 1.21 kg m**□** ³), improving pore structure and enhancing water and oxygen infiltration. Water saturation cycles (30–70% MWHC) further promoted oxygen transport within the BR matrix.

### Microbial community shifts associated with sulfur amendment

Microbial community composition in G_OM and G_OM3S was profiled at days 58, 155, and 282 using SSU rRNA amplicon sequencing, and showed clear separation between treatments, with S_8_ amendment the major driver of divergence (**Fig. 2a**); composition was stable between days 155 and 282, suggesting adaptation to the evolving geochemistry. While the two treatments shared several dominant taxa, the genus *Thioalkalivibrio*_A, a likely alkaliphilic sulfur oxidizer^26^, was up to ∼5-fold more abundant in G_OM3S (up to relative abundance 6.87%) than in G_OM (relative abundance g 0.04%) by day 155, coinciding with sulfate production (**Fig. 2b**, **Fig. 1e** and **f**, **Supplementary Fig. 4** and **Supplementary Tables 4** and **7**). These patterns are consistent with a shift towards microbial communities associated with sulfur-rich conditions.

**Figure 2:**
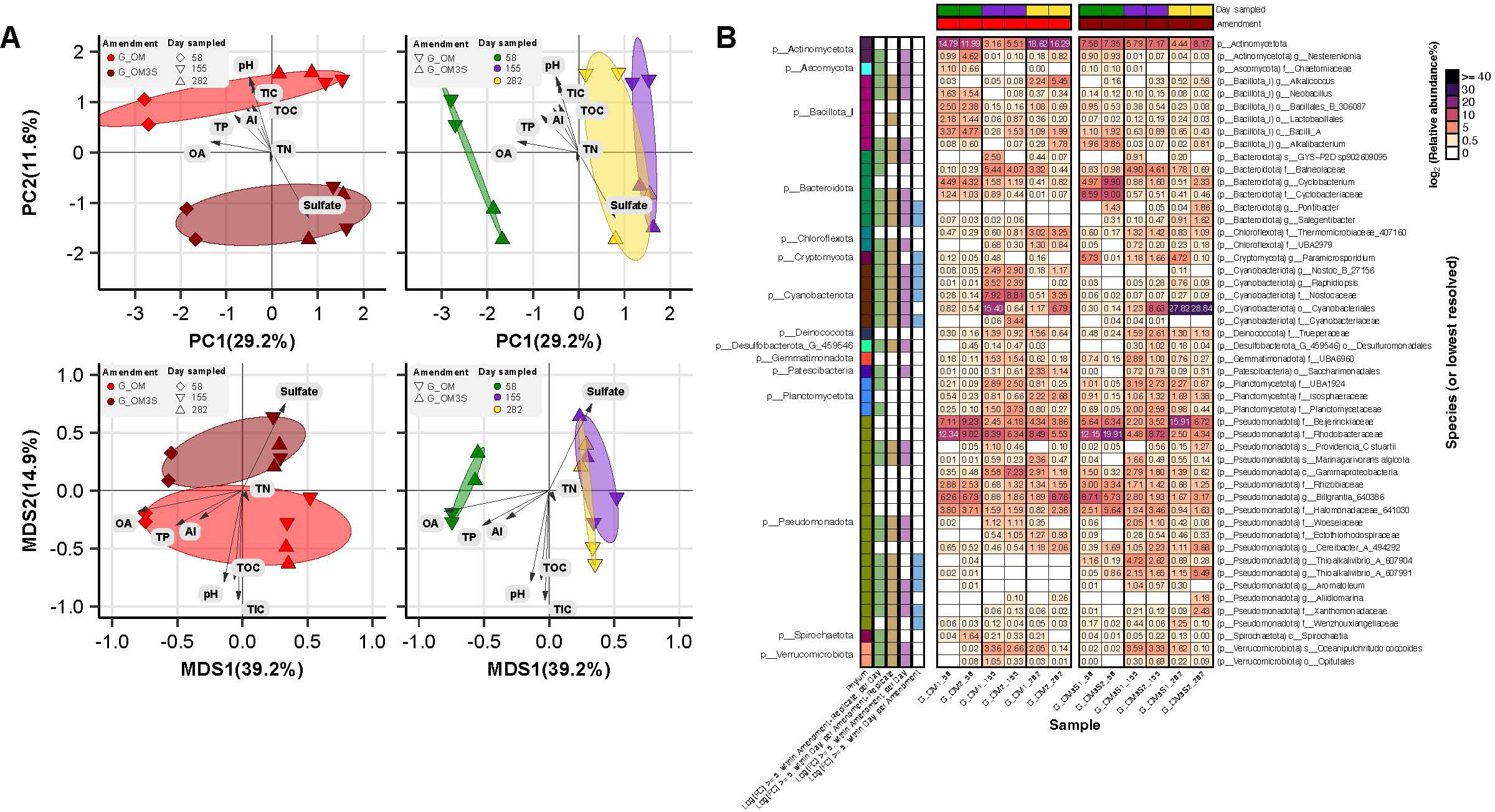
a) PCA of the Euclidean distances for robust centred log-ratio (rclr) values, and ordination analysis of Jaccard dissimilarities (MDS), visualizing the variation in microbial community composition across the glasshouse trial control (G_OM) and sulfur-amended (G_OM_3S) treatment groups at different time points. Environmental variables fitted to the ordinations using envfit are annotated with arrows and scaled to aid visualization. **b)** Heatmap of the rRNA relative abundances of microbial taxa across G_OM and G_OM_3S groups at different time points. Lineages that had a log-fold change ≥ 5, between or within amendments, day of sampling, and/or replicate, according to edgeR have been annotated on the left-hand side. Taxonomic ranks for each lineage are indicated by the prefixes p (phylum), c (class), o (order), f (family), g (genus). A suffix in a genus or species name, e.g. ‘_A’ in *Thioalkalivibrio*_A, is used in genome taxonomy database (GTDB) annotations to indicate polyphyletic groups, or to indicate the group has been subdivided based on taxonomic rank normalization according to the current GTDB reference tree. In the GreenGenes2 database, polyphyletic groups are indicated by a numeric suffix (e.g., ‘_607904’).

### Metabolic potential and expression of sulfur-cycling pathways

Metagenomic and metatranscriptomic sequencing of bulk G_OM3S samples collected at days 155 and 282 was used to identify sulfur transformation pathways and their expression, capturing community-level activity within a heterogeneous matrix with redox gradients. Sequencing produced 15.9 Gbp of metagenomic and 4.3 Gbp of transcriptomic reads, respectively, enabling assessment of both functional potential and *in situ* gene expression. From the metagenomic data SingleM estimated 75% of the reads to be microbial and Nonpareil reported 81.9% community coverage, indicating that a substantial portion of the microbial community was sequenced (**Supplementary Fig. 5**, **Supplementary Tables 8** and **9**). Assembly and binning yielded 254 genomic bins representing *in situ* microbial populations (**Supplementary Table 6**), of which 71 met quality thresholds for metagenome-assembled genomes (MAGs; quality **≥** 50, defined as completeness − 3 × contamination). Approximately 39% of the metagenomic reads mapped to these 71 MAGs (54.2% to all bins), indicating that a substantial fraction of the community is represented by these genomes (**Supplementary Table 5**). The MAGs span 14 bacterial phyla and broadly recapitulate the diversity observed by SSU rRNA amplicon sequencing (**Supplementary Fig. 4**). Many remained unclassified at lower taxonomic ranks, suggesting the presence of previously undescribed microorganisms (**Supplementary Fig. 6**). Optimal growth conditions predicted by GenomeSpot^27^ indicated that all MAGs are derived from haloalkalitolerant organisms (**Supplementary Fig. 7**).

Metabolic reconstruction of the MAGs using DRAM^28^, HMSS2^29, 30^ and additional HMM-based analysis (**Supplementary Note 2**), revealed extensive capacity for coupled carbon and sulfur cycling (**Fig. 3**, **Supplementary Fig. 8**, **Supplementary Tables 10**–**16** and **Supplementary Note 3**). Genes for core carbon metabolism, polysaccharide degradation, and sulfur cycling were widespread, supporting active organic matter turnover and sulfur transformation potential. Initial HMSS2 screening identified putative polysulfide, thiosulphate and sulfur reductase (Psr/Phs/Sre-like) catalytic subunit homologs across 38 MAGs, although co-encoded subunits (PsrB/PsrC-like) were sparse, with complete three-subunit architecture recovered in only one MAG (**Fig. 3**, **Supplementary Fig. 8**). Because catalytic subunits within the Mo/W-bisPGD binding (MopB) superfamily are functionally diverse and difficult to distinguish from one another using sequence similarity alone^31, 32^, a targeted classification approach integrating phylogenetic placement, transmembrane topology prediction and gene-neighborhood analysis was applied (**Supplementary Note 2**). This identified PsrA/PhsA/SrrA-clade homologs in two genomes – JARFUG01 bin58 (order *Desulfuromonadales*) and the lower-quality CSSed10-48 bin154 (family *Trueperaceae*) – with neighboring subunit homologs consistent with PsrABC/PhsABC-like reductase architectures (**Supplementary Figs. 9** and **10**, **Supplementary Table 17**). Both loci also encoded an adjacent rhodanese-type sulfurtransferase, consistent with polysulfide trafficking roles described in model sulfur respirers^33^. However, definitive assignment of canonical polysulphide reductase activity was precluded by the close evolutionary relationships of MopB homologs and incomplete subunit recovery. Sulfur oxidation potential was broadly distributed, with pathways including *sqr* (sulfide:quinone oxidoreductase), *sox* (sulfur oxidation complex), *shdrABC* (heterodisulfide reductase, potentially operating in reverse in sulfur oxidation pathways), *soeABC* (sulfite:quinone oxidoreductase) and *dsrABC*–*aprAB*–*sat* (dissimilatory sulfite reductase–adenylylsulfate reductase–sulfate adenylyltransferase), supporting the oxidation of sulfide, sulfane sulfur and thiosulfate to sulfate.

**Figure 3:**
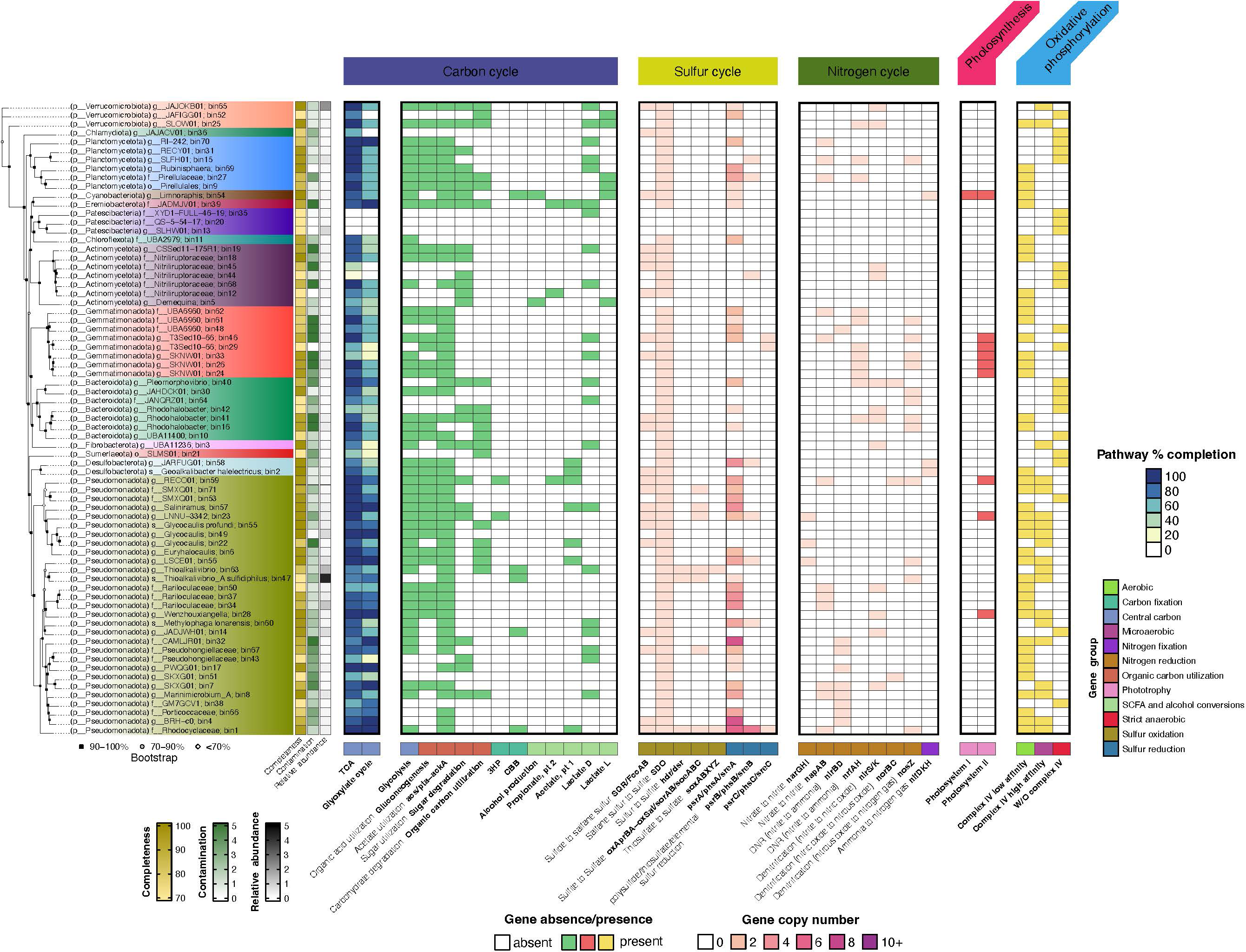
Heatmap illustrates the metabolic traits of the 71 higher-quality MAGs and corresponding phylogenetic tree. The tree leaf label for each genome has been colored by the phylum classification. The metabolic pathways are categorized into five main metabolism types: Carbon, Sulfur, and Nitrogen cycle, and Photosynthesis and Oxidative phosphorylation.

Expression data indicate *in situ* activity of sulfur and carbon cycling genes across redox gradients (**Fig. 4** and **Supplementary Table 18**). Transcripts from several PsrA/PhsA/SrrA-clade MopB candidates were detected, with the strongest signal in JARFUG01 bin58, where the catalytic subunit, neighboring PsrB-like, and downstream rhodanese-type sulfurtransferase reached 112, 96 and 134 TPM, respectively. Expression of the catalytic and B-like subunits was also detected in the lower-quality CSSed10-48 bin154 (36 and 37 TPM respectively), with the downstream rhodanese-like sulfurtransferase expressed at 243 TPM supporting a potential role in polysulfide reduction. Sulfur oxidation activity was primarily associated with *Thioalkalivibrio* species. *Thioalkalivibrio* bin63 encodes and expressed the truncated *sox* system and *shdrABC* and *soeABC* (TPM 19–1143), which facilitate the oxidation of thiosulfate and sulfane sulfur to sulfate. *Thioalkalivibrio sulfidiphilus* bin47 expressed *dsrABC–aprAB–sat* and *sHdrABC–aprAB–sat* (TPM 38–375), indicating its potential role in sulfane sulfur oxidation to sulfate, together with expressed *sqr* V and *sdo* I (TPM 21 and 31), indicating its capacity to oxidize sulfide to sulfite via glutathione-bound intermediates, with sulfite subsequently contributing to sulfate production or thiosulfate formation. Expression of cbb**□**-type oxidases in *Thioalkalivibrio* bin63 and *Thioalkalivibrio sulfidiphilus* bin47 indicates adaptation to redox gradients, while co-expression of *acs*, *ldhA*, and Calvin cycle genes (TPM **≥** 17) suggests a potential mixotrophic lifestyle. In addition, expression of *sqr* and *sdo* homologs in *Limnoraphis* bin54 and *Aquisalimonadaceae* bin56 suggests sulfide oxidation potential beyond canonical sulfur-oxidizing taxa. *Limnoraphis* bin54 also expressed genes for photosystems I and II, *nifHDK*, and glycoside hydrolases (enzymes involved in the degradation of plant-derived polysaccharides), indicating a phototrophic and metabolically versatile lifestyle capable of utilizing organic substrates.

**Figure 4:**
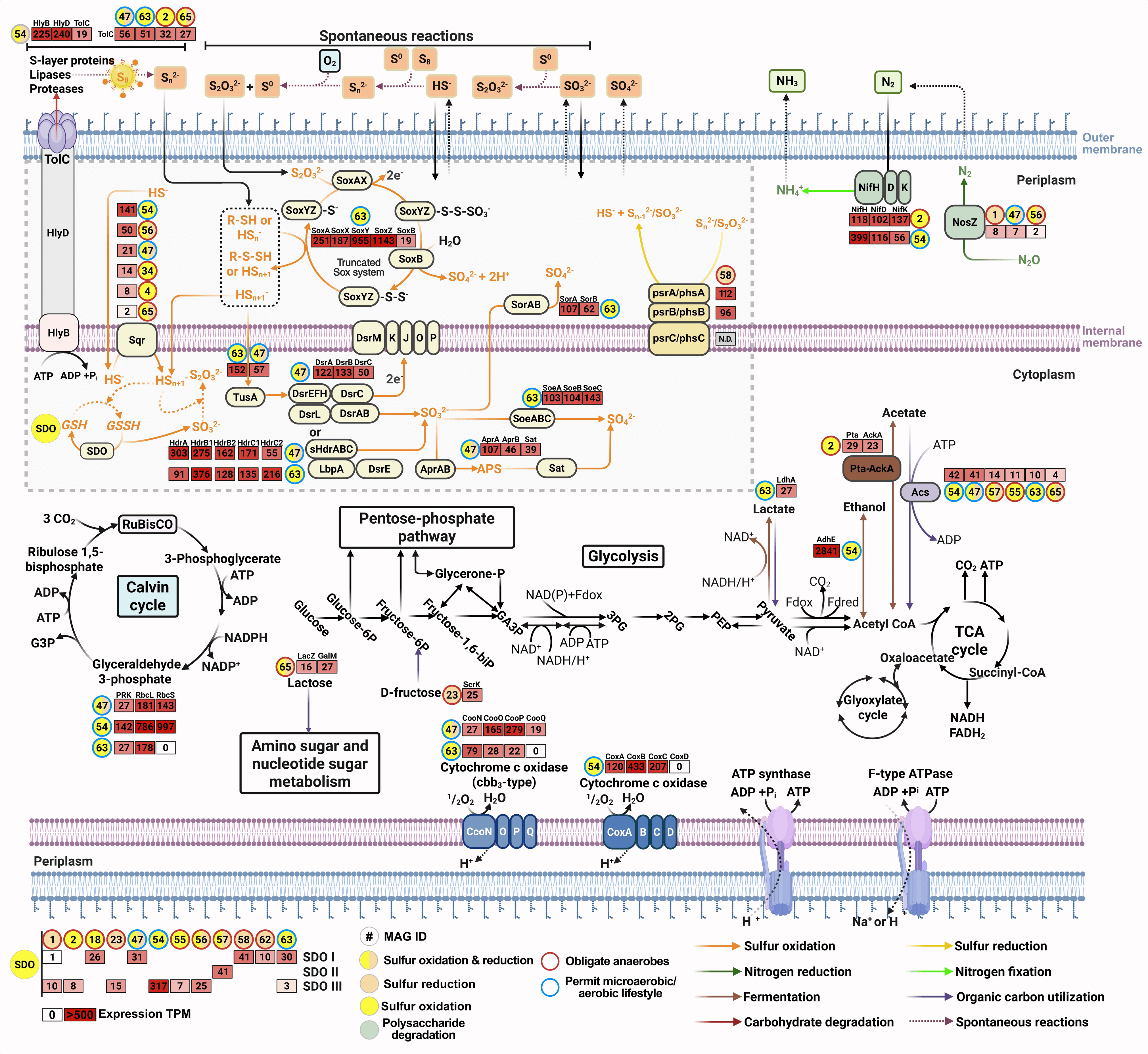
Inferred metabolic gene network of the major microbial taxa associate with carbon, sulfur and nitrogen metabolism. MAGs encoding each gene are indicated by numbered colored circles. The transcripts per million (TPM) expression levels of each gene are indicated by the numbered white-to-red gradient-colored boxes. Created in part with BioRender.com.

Genomic and transcriptomic data collectively suggest sulfur transformation potential in the G_OM3S system is mediated by a distributed microbial network, encompassing both oxidative and reductive pathways. These predictions were tested in controlled batch experiments.

### Redox-controlled batch experiments resolve the microbial–abiotic relay

Batch incubations of G_OM3S-derived communities (pH 10.3, 0.6 M Na□) isolated the individual sulfur transformations under controlled oxygen and carbon supply: S_8_ alone (B_S), + inoculum (B_SI), + cellulose (B_SC), and + cellulose + inoculum (B_SCI), in open-air flasks (spatial redox gradients) or sealed serum bottles (anaerobic) (**Fig. 5a** and **b**).

**Figure 5:**
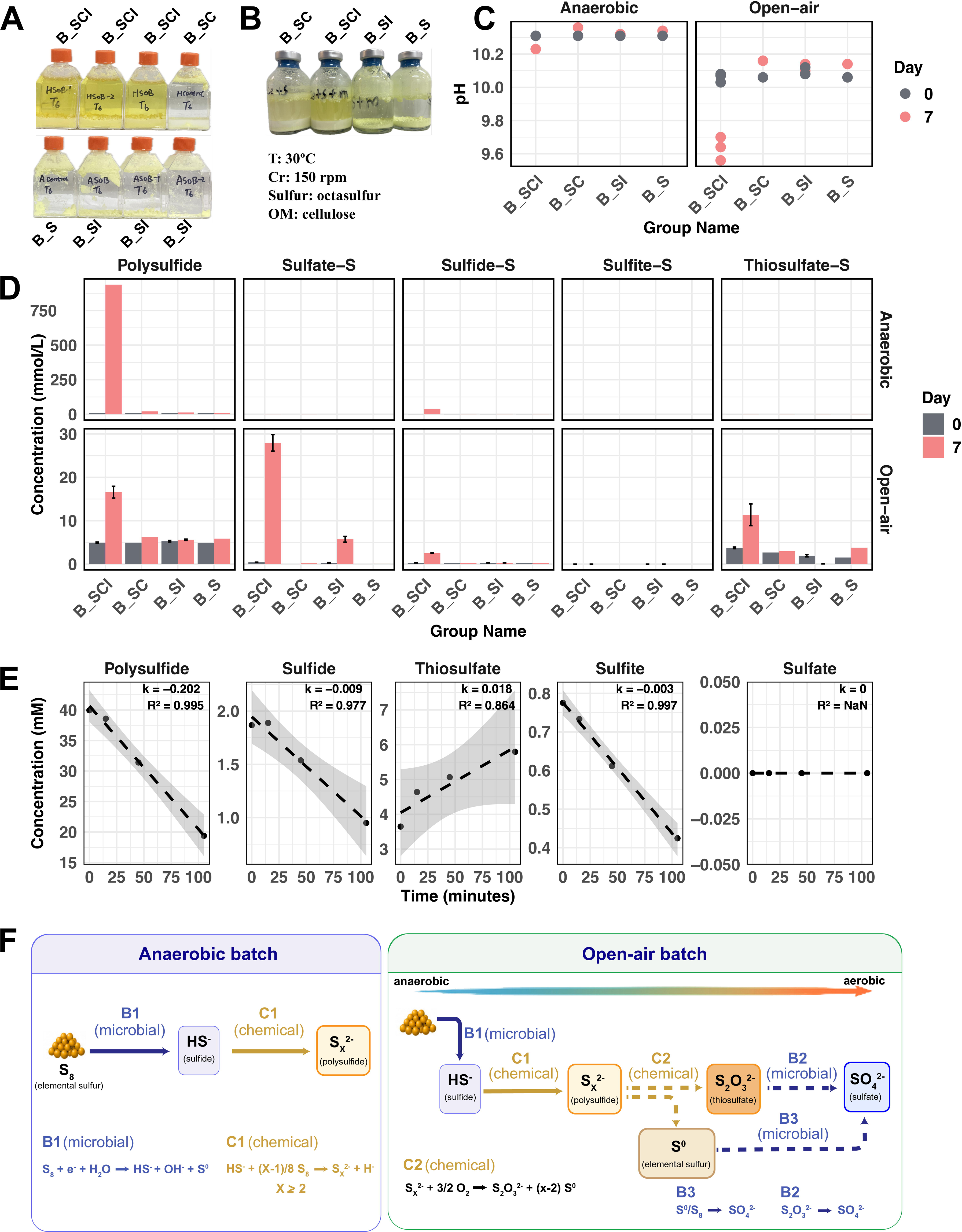
**(a)** Open-air flasks with oxygen flux and **(b)** anaerobic conditions with sealed serum bottles were used. The experiment groups include different substrates: B_SC (S_8_ + cellulose), B_SCI (S_8_ + cellulose + inoculum), B_S (S_8_), and B_SI (S_8_ + inoculum). **(c)** Dynamics of pH after one week of incubation in open-air and anaerobic batches. **(d)** Concentration of sulfur species: polysulfide, sulfate-S, sulfide-S, sulfite-S, and thiosulfate-S (mean ± s.d., n = 3) **(e)** Chemical oxidation rate of the polysulfide liquid at room temperature in open-air conditions (k values: rates of change in concentration per unit time and R²: coefficients of determination). **(f)** Schematic illustration of sulfur transformations in batch incubations with elemental sulfur and organic matter under (left) anaerobic and (right) open-air conditions. Major microbial (B1–B3) and abiotic (C1–C2) pathways are indicated. Solid arrows indicate reactions under oxygen-limited conditions; dashed arrows indicate reactions occurring upon oxygen exposure.

Under anaerobic conditions, only B_SCI produced substantial polysulfide and sulfide after 7 days (**Fig. 5d**). These results indicate that the reduction of S_8_ to sulfide (**Fig. 5f**, **reaction B1**) is microbially mediated and dependent on organic substrates. The sulfide produced under alkaline conditions then abiotically reacts with S_8_ to form polysulfides (**Fig. 5f**, **reaction C1**)^34–36^, consistent with the greater accumulation of polysulfide than sulfide. A cell-free chemical assay confirmed that polysulfide oxidizes to thiosulfate rapidly on oxygen exposure at pH 9.6 (0.018 mM min**□** ¹; **Fig. 5e**, **reaction C2**) but produced no sulfate demonstrating that the acid-generating terminal oxidation to sulfate is exclusively microbial (**Fig. 5f**, **reactions B2–B4**).

Under open-air conditions, only B_SCI reduced solution pH and accumulated soluble sulfur (49.08 ± 1.18 mM), including reduced species (polysulfide, 11.66 ± 2.21 mM; sulfide, 2.25 ± 0.17 mM) and oxidized species (sulfate, 27.54 ± 3.24 mM; thiosulfate, 7.62 ± 4.66 mM), whereas controls showed negligible transformation and pH change (**Fig. 5d**). The coexistence of reduced and oxidized sulfur species indicates simultaneous anaerobic and aerobic processes operating in spatially separated redox zones (**Fig. 5f**). In oxygen-limited zones, microorganisms reduced S**□** to sulfide using organics (**Fig. 5f**, **reaction B1**), which then reacted abiotically with S**□** to produce polysulfides (**Fig. 5f**, **reaction C1**), consistent with the anaerobic B_SCI cultures. In microaerobic and aerobic zones, polysulfides were abiotically oxidized to thiosulfate and reactive elemental sulfur (S**□**) (**Fig. 5f**, **reaction C2**). Microorganisms then oxidized sulfur intermediates, potentially including thiosulfate, sulfane sulfur, and sulfide, to sulfate (**Fig. 5f**, **reactions B2–B4**). Organic matter strongly amplified the cycle: relative to B_SI with limited S_8_ transformation, B_SCI contained >10-fold more total organic carbon and organic acids, with 12-fold higher sulfur transformation and 6-fold higher sulfate production (**Supplementary Fig. 11**). This intensified sulfur cycling underpins the pH decrease observed only in the open-air B_SCI treatment (**Fig. 5c**). Finally, rRNA amplicon profiling of B_SCI biomass revealed distinct microbial communities between anaerobic and open-air batches. The anaerobic batch culture was dominated by a distinct set of taxa, while open-air cultures showed greater diversity, encompassing the taxa found in the anaerobic batch alongside additional taxa (**Supplementary Fig. 12**). Together, these results support a self-amplifying microbial–abiotic sulfur relay operating across spatially separated redox zones, accelerating the conversion of S**□** into a reactive sulfur pool that microbial communities oxidize to sulfuric acid for dealkalization.

### Field application of the microbial-abiotic sulfur relay for BR dealkalization

An ongoing, large-scale field trial (∼1 hectare) was set up on the Queensland Alumina Ltd site using seawater-carried (i.e., seawater-treated) BR of strong alkalinity (pH 9.5–10) and extreme salinity (EC 70–100 mS/cm). Over 687 days, the dual-amendment treatment combining 40% OM (volume to volume) and 3% S_8_ (weight by weight) (F_OM3S) drove rapid and sustained dealkalization relative to OM alone (F_OM): porewater pH declined from 9.47 to 8.72 ± 0.41, whereas F_OM rebounded to 9.79 ± 0.05 (**Fig. 6a**). These findings confirm that the OM-S_8_ strategy enhances and accelerates BR dealkalization under field conditions, consistent with our findings in the laboratory and glasshouse experiments. In the F_OM3S plot, the resulting pH conditions supported spontaneous colonization by local plant species, with dense grass cover consistent with early technosol formation (**Fig. 6b**).

**Figure 6:**
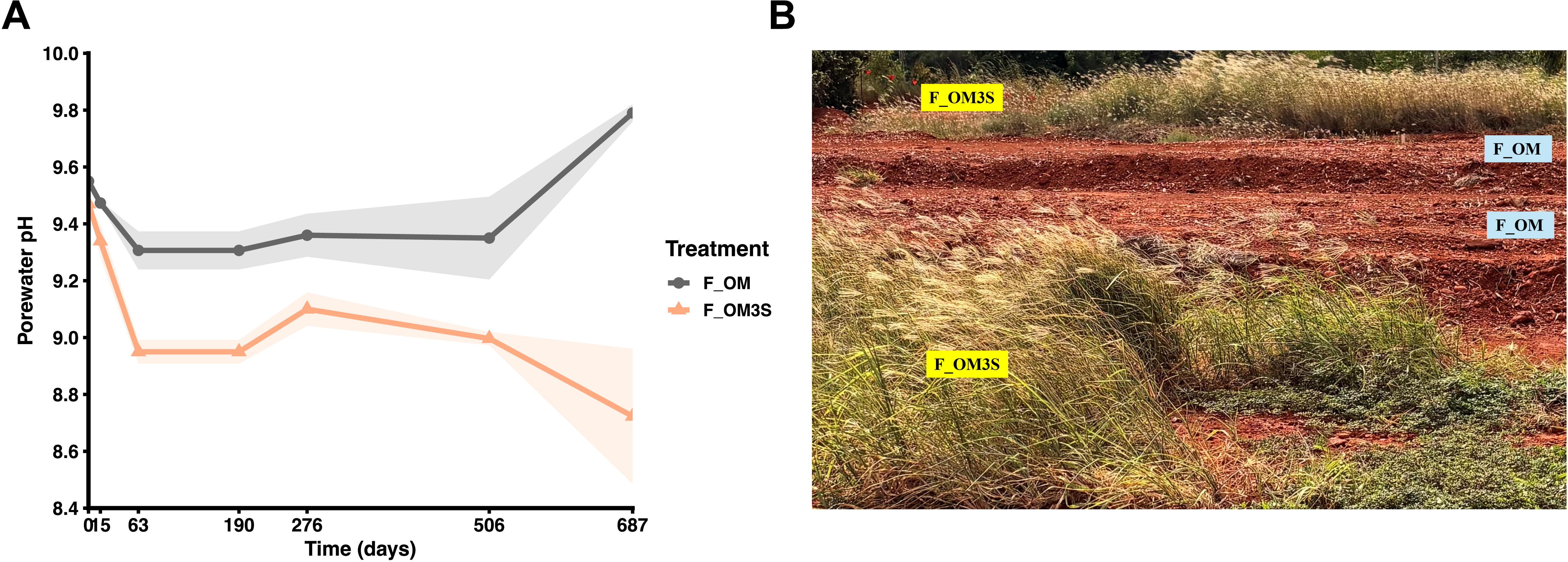
Field experiment results for soil pH and plant colonization. **(a)** Dynamics (i.e., a time-series of plot-specific monitoring) of bauxite residue pH from May 2023 to August 2025 for two treatments: F_OM (40% v/v plant mulch) and F_OM3S (40% v/v plant mulch supplemented with 3% elemental sulfur). **(b)** Photograph taken in August 2025 showing pioneer plant colonization only in the F_OM3S treatment.

## Discussion

This study establishes a self-amplifying microbial–abiotic sulfur relay that enables rapid, sustained *in situ* acid generation and dealkalization of bauxite residue (BR). This mechanism was demonstrated across laboratory and glasshouse experiments, and a one-hectare field trial, where a pioneer plant community developed. The relay is activated by co-amendment with organic matter and elemental sulfur (S_8_), and overcomes two barriers that impede sulfur-driven biological treatment of BR: the near-complete insolubility of S_8_ ^37^, and extreme haloalkaline conditions (pH >10, EC >40 mS cm**□** ¹) that severely constrain microbial activity^38, 39^.

A central feature of the relay, demonstrated by redox-controlled batch experiments, is the reinforcing interaction between microbial and abiotic sulfur transformations. This coupling provides the mechanistic basis for dealkalization in OM–S_8_ amended BR and represents a deliberately engineered remediation strategy not previously described for haloalkaline systems. The relay bypasses the constraint of S_8_ insolubility through a chemical amplification step: sulfur-reducing microorganisms initiate S_8_ turnover using organic carbon as an electron donor under oxygen-limited conditions, producing HS^—^ pulse that reacts abiotically with solid S_8_ to generate soluble 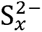. This single abiotic reaction amplifies the products of microbial sulfur reduction into a disproportionately large and diffusible pool of reactive intermediates, the key step that makes the system self-amplifying. In anoxic microsites, these intermediates are microbially reduced back to HS^—^, which dissolves additional S_8_. Upon diffusing into oxic zones, 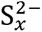 undergoes abiotic transformation to thiosulfate and reactive S**□** ^40, 41^, expanding the diversity of bioavailable sulfur species alongside the HS^—^ and 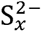 already present. Each of these intermediates represents a potential substrate for microbial oxidation, enabling multiple oxidative pathways to operate in parallel^42^. Critically, the terminal oxidation of soluble sulfur intermediates e.g., thiosulfate to sulfate (the acid-generating step that drives dealkalization) is exclusively microbially mediated, as sulfate formation was detected only in treatments where both oxygen and active microbial communities were present. This confirms that chemical transformations amplify sulfur bioavailability, but microbial activity remains essential for net acidification.

Metagenomic and metatranscriptomic analyses revealed a phylogenetically diverse microbial network capable of sustaining the coupled oxidative and reductive sulfur transformations central to the relay. The oxidative arm is well resolved: *Thioalkalivibrio* spp. were the dominant sulfur oxidizers, with bin63 expressing *sox* and *sHdrABC*–*soeABC*, and *T. sulfidiphilus* bin47 expressing *dsrABC*–*aprAB*–*sat* together with *sqr* and *sdo*, consistent with the capacity to oxidize thiosulfate, sulfane sulfur, and sulfide to sulfate. Sulfide oxidation potential also extended to *Limnoraphis* bin54 and *Aquisalimonadaceae* bin56, suggesting that the oxidative network is broader than canonical sulfur-oxidizing taxa alone. The reductive arm presents a more complex picture. The co-expression of rhodanese-like sulfurtransferases alongside catalytic subunits in JARFUG01 bin58 (order *Desulfuromonadales*) suggests that polysulfide trafficking may be integral to this process, consistent with known roles of rhodanese proteins in model sulfur respirers^33^. Whether these organisms encode functionally novel reductase architectures or use alternative sulfur-reduction pathways cannot be resolved by metagenomic inference alone and remains an open question. Definitive resolution will require culture-based characterization or stable isotope probing (SIP) to directly link phylogenetic identity with functional activity. Together, the combined genomic, transcriptomic, and experimental evidence supports a community-level division of labor in which oxidative and reductive sulfur cycling are partitioned across different taxa adapted to the contrasting redox microenvironments generated within the BR matrix.

This microbial–abiotic architecture delivers a remediation outcome that prior approaches have been unable to achieve. Fermentative organic acids are much weaker than mineral acids such as sulfuric acid, and cannot remove the recalcitrant alkali (Na) bound within aluminosilicate minerals; gypsum amendment stabilizes pH at 9–10 without addressing solid-phase alkalinity^15, 16^; and direct treatment with mineral acids is financially and environmentally prohibitive at industry scale. The relay overcomes these constraints through the *in situ* generation of sulfuric and organic acids from low-cost, agricultural-grade inputs, lowering porewater pH sufficiently to substantially lower metal(loid) solubility and improve physical and chemical conditions for plant community growth and development at field scale. This outcome transforms BR from a persistent liability requiring engineered containment into a substrate capable of supporting ecological recovery. More broadly, coupling biotic reductive initiation and abiotic chemical amplification with biotic terminal oxidation may provide a general design principle for remediating extreme haloalkaline industrial wastes.

## Material and methods

### Experimental design and set up

#### Laboratory proof-of-concept

Lab-scale (L_) experiments were conducted to test the feasibility of microbial sulfuric acid production from agricultural-grade elemental sulfur (S_8_) in extreme haloalkaliphilic bauxite residue (BR) and the potential acceleration of S_8_ oxidation by organic matter. These experiments assessed whether S**□** could be biologically oxidized to sulfuric acid under these conditions to facilitate BR dealkalization. Alkaline BR were sourced from the residue disposal area at the Rio Tinto Gove refinery (Northern Territory, Australia) in 2023, which was air-dried in an oven (40–45 °C) and ground to pass a 2 mm sieve. The organic matter comprised aged plant mulch derived from local vegetation cleared during mining operations at the Rio Tinto Gove refinery, was processed by ballmill to a fine particle size (< 2mm). Four treatment groups were set up: L_BR: BR without amendments as a control, L_3S: BR + S_8_ (3% w/w; 3S), L_OM: BR + OM (i.e., 40% v/v ground plant mulch; OM), and L_OM3S: BR + OM + 3S (OM3S) (**Supplementary Table 1**). The mixtures of BR and amendment(s) were placed in 200 ml containers, which were capped, mixed thoroughly, and allowed to settle for 15 minutes before removing the cap. The dry mix was watered to saturation (i.e., 100% water holding capacity) before commencing incubation. The amount of water required to reach saturation in each treatment was estimated from a pre-saturation test ^43^. A Rhizon porewater sampler was inserted into the center of each container, which was then watered with deionized water to approximately 100% maximal water-holding capacity (MWHC) to commence incubation. The incubation was carried out in a Biora RIC600 incubator under controlled conditions: 30°C temperature, ∼55% relative humidity (RH), and 20% ambient light brightness. During the incubation, the container experienced wet-dry cycles with water contents of approximately 30–70% MWHC. Porewater samples in each container collected at different time points for monitoring chemical changes, including pH, EC, sulfate concentration, total oxidative sulfur, total phosphorus (TP), total organic carbon and organic acid.

#### Glasshouse trial

The glasshouse trial (G_) used bauxite residue (BR) bulk-sourced from a storage pond located at Rio Tinto Gove refinery (Nhulunbuy, Arnhem Land in the Northern Territory of Australia) and coarse tree mulch (approximately 30mm in length, sourced from local trees removed for mining activities). The trial was conducted between September 2022 and May 2023 at St Lucia campus, University of Queensland (Brisbane, QLD, Australia). The ambient temperature in the glasshouse ranged from 10 to 30°C (**Supplementary Fig. 1**), resembling uncontrolled field conditions. The amount of mulch added into the BR was 40% on volume by volume (v/v) basis (equivalent to about 5% on weight basis). Bulk density of BR with and without mulch was measured *in situ* using stainless steel rings (5.6 cm diameter × 4 cm height). Sample cores were collected, oven-dried at 105 °C until constant weight and then weighed to determine density. Elemental sulfur (S_8_, agricultural grade) was subsequently admixed into the mixture of BR and mulch on a weight-by-weight basis (**Supplementary Table 2**). The treatment groups tested in this trial include combinations of three factors - BR, OM and S_8_ : G_OM: BR + OM (40% v/v), G_OM0.5S: BR + OM0.5 (0.5% S_8_, w/w), G_OM1S: BR + OM1S (1% S_8_ w/w), and G_OM3S: BR + OM3S (3% S_8_ w/w).

The blended mixtures were transferred into pots without cover lids, with two replicates per treatment. A Rhizon sampler (MOM, Rhizosphere Research Products, Wageningen, The Netherlands) for *in situ* sampling porewater was inserted into the center of each pot, in 24 hours after the pots were watered with deionized water to approximately 100% maximal water-holding capacity (MWHC). The incubation of the pots commenced with the saturated watering in a glasshouse. During the incubation, the pots experienced cyclic wet/dry intervals of approximately 30–70% MWHC.

Aliquots of 10ml porewater were sampled from each pot at multiple intervals, of which subsamples of 5 ml were immediately frozen for the analysis of: total oxidative sulfur (TOS), total phosphorus (TP), Al, Ca, Mg, Si, Fe, and Na, total organic carbon (TOC) and organic acid, and sulfate. The rest of the fresh porewater samples were immediately used for measuring pH and EC. The solid samples were collected at days 58, 155, 282 for DNA extraction and sequencing. At the end of the experiment, the solids were sampled for geochemical and mineralogical analyses: including pH_1:5_ as pH measured in water extracts (solid/water = 1:5 (w/w)), the alkalinity of tailings, and cation exchange capacity (CEC), qXRD, solid TOC and total inorganic carbon (TIC).

#### Microbial sulfur transformation and sulfate production assay

To determine the factors regulating sulfate production and the associated pathways, we conducted a batch experiment (B_) using mixed BR samples from the G_OM3S treatment (days 155 and 282 of the glasshouse trial) as the inoculum. The mixed solid samples were cultured in a 100 ml bottle containing a medium similar to the porewater of the glasshouse trial, with a pH of 10.3 and 0.6M sodium. Key factors considered included the availability of microorganisms, organic matter, and oxygen.

The experimental groups were the control S**□** only (B_S), S_8_ + inoculum (B_SI), S_8_ + cellulose (B_SC), S_8_ + cellulose + inoculum (B_SCI) (**Supplementary Table 3**). Oxygen availability was tested in these groups using open-air flasks to simulate microaerobic conditions with both oxic and anoxic zones, and sealed serum bottles to create anaerobic environments. To investigate the chemical oxidation of polysulfide, after 7 days of incubation, 100 ml medium was collected and filtered through a 0.22 **μ**m Millex-GP syringe filter (SLGPR33RS, Merck) into a 250ml flask and left in a fume hood. Cellulose was used as a proxy of organic matter since plant mulch contains abundant cellulosic carbohydrate. Monitored sulfur species included polysulfide, sulfide, thiosulfate, sulfite, and sulfate, along with pH. Soluble sulfur species (sulfide, thiosulfate, sulfite, and sulfate) were quantified using a Dionex ICS-2000 ion chromatography (IC) system equipped with an AS50 autosampler. A 25 µL sample injection volume was used for all analyses. Separation was achieved on a Dionex IonPac AS18 analytical column (4 × 250 mm; PN 060549) coupled with a Dionex IonPac AG18 guard column (4 × 50 mm; PN 060551) at 35 °C. The eluent was generated using a Dionex EGC III KOH cartridge (PN 074532). The elution program consisted of an initial concentration of 12 mM KOH, ramping to 34 mM over 5 min with a 3 min hold, followed by a ramp to 55 mM over 1 min and a 5.5 min hold, for a total run time of 19.5 min. A Dionex AERS 500 suppressor (4 mm; PN 082540) was used for conductivity suppression. Detection was performed using a Dionex DS6 heated conductivity cell (35 °C) in combination with a Dionex variable wavelength detector (VWD) set at 230 nm. Quality control was ensured using a CertiPur Anion Multi-element Standard II (lot HC31103348; Merck), diluted 5× with Milli-Q water. The diluted standard contained chloride (199.80 mg/L) and sulfate (66.66 mg/L). Sample and standard preparation followed published methods ^44^. Polysulfide concentration was calculated by the sulfide–polysulfide equilibrium (Eq. (1)). In the presence of excess amounts of elemental sulfur, the average polysulfide chain length, *x* =4.91 (21 °C) and the equilibrium can be represented by ^35, 45, 46^:

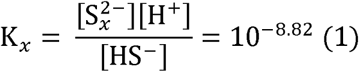

#### Field trial

An ongoing, long-term field trial (F_) was established in the BR storage site of Queensland Alumina Ltd (Gladstone, Queensland) from May 2023, covering approximately 1 hectare (60**□** m × 150**□** m, including control and peripheral margin area), to evaluate the sulfur-cycle strategy in seawater treated (i.e., carried in seawater) BR (ST-BR) exhibiting strongly alkaline pH (9.5–10) and extremely saline (saturated EC 70–100 mS/cm) conditions. The large plots (i.e., main plots) (length X width X depth - 3 m X 95 m X 0.7 m) were set up using air-dried and crushed ST-BR, overlaying the sublayer of compacted ST-BR. The neighboring main plots were separated by a 0.5**□** m wide interrow to accommodate irrigation infrastructure, operational access, and lateral leaching process in response to irrigation and/or rainfalls in wet seasons. The plots all received a basal amendment that included 40% (v/v) plant mulch, which was admixed mechanically into the profile. In combination with the above basal additions, the treatment of F_OM3S was applied with a designated rate (% w/w air-dry BR) of 3% of agricultural-grade elemental sulfur (S_8_) (equivalent to 5024 kg/ha for remediating 20 cm layer). Agricultural machinery was used for spreading and admixing both mulch and elemental sulfur into BR profiles throughout the ∼70 cm depth of the bauxite residue profile. To characterize baseline geochemical conditions—samples were collected from the top 10**□**cm of treatment plots prior to amendment. To assess geochemical dynamics after amendment, treatment samples were collected on 30/05/2023 (Day 0), 13/06/2023 (Day 14), 1/08/2023 (Day 63), 6/12/2023 (Day 190), 17/10/2024 (Day 506), 16/04/2025 (Day 687). Samples were taken from the top 10**□** cm of each subplot, which were placed in plastic bags and immediately stored in pre-cooled insulated containers with ice packs. Samples were transported at 4**□** °C to the laboratory. Upon arrival, aliquots of subsamples were oven-dried at 40**□** °C until constant weight. The dried material was gently ground with a mortar and pestle and sieved through a 2**□** mm mesh. The < 2**□** mm fraction was used for porewater pH analyses.

### Mineralogical and geochemical analysis

#### Porewater

The pH and EC in porewater were measured using a pH electrode (HORIBA) and an EC electrode (TPS 2100), respectively. The TOC and TN were quantified using a TOC analyzer with a TN detector (VCPH, Shimadzu, Japan). Aliquots of porewater samples were acidified and diluted with 2% nitric acid for the analysis of elements (i.e., K, Na, Ca, Mg, Si, Al, Fe) by means of Inductively Coupled Plasma Optical Emission Spectroscopy (ICP-OES) (Varian Vista Liberty, Australia). Sulfate concentrations were measured by means of ion chromatography (IC, DX-120, Thermo Scientific). Concentrations of 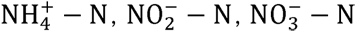 and 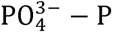 were quantified using a Lachat QuikChem8000 Flow Injection Analyzer (Lachat Instrument, USA).

#### Solid samples

Solid samples collected at the end of the glasshouse trial were used for the analysis of pH, CEC, qXRD, TOC and TIC, and residual alkalinity. The pH and electrical conductivity (EC) in 1:5 water extracts were measured by using a pH electrode (TPS 900) and an EC electrode (TPS 2100). Total nitrogen (TN) contents were determined by a LECO CNS-2000 Analyzer (LECO Corporation, MI, USA). For TOC, inorganic carbon in the samples were removed by HCl pretreatment and the TOC contents were determined by a LECO CNS-2000 Analyzer. The measurement of CEC was modified from Section 15A1 in (Rayment and Lyons, 2011) and (Liu et al., 2007). Duplicates of 2g aliquots were extracted with 40ml milliQ H_2_O and 1M NH_3_Cl (adjusted to pH 7), respectively, on end-on-end shaker for 1hour. The extracts were centrifuged (at 3220xg) for 10 mins and filtered through Whatman #42 filter paper. The filtered extracts were diluted with 2% HNO_3_ for the analysis of major cations by using an ICP-OES. The alkalinity was measured according to the modified Sobek ANC test^47^ (**Supplementary Note 1**).

The mineralogy of samples was characterized by X-ray Powder Diffraction (XRD) and quantitative XRD (q-XRD). The XRD spectra of the samples were collected from 2**θ** = 2° to 80° with 0.02° per step using a D8 Advanced Diffractometer (Bruker AXS) with a LynxEye detector. The q-XRD was conducted to estimate the composition of mineral phases in the samples by mixing fluorite (CaF2) as an internal standard, at a 10% (w/w) ratio. Fluorite was selected as its diffraction pattern does not significantly overlap with the patterns of the specimen, and it has good stability with the materials. Quantitative Dominant minerals were identified from the spectrum using the Diffracplus Evaluation Package V5.1 (Bruker AXS, Germany) based on the PDF-4 mineral database (2020 release). The quantification of the identified minerals was performed using TOPAS V6.

### rRNA amplicon sequencing and analysis

#### DNA extraction and PCR amplification

Microbial DNA was extracted from the glasshouse samples collected at days 58, 155, and 282, by using a DNeasy PowerSoil Kit (Qiagen), and the concentration and quality evaluated using a Nanodrop ND-2000 (Thermo Scientific, US). The SSU ribosomal RNA (rRNA) genes in the extracted DNA were then PCR amplified with the primers 926F (5**′**-AAACTYAAAKGAATTGACGG-3**′**) and 1392R (5**′** ACGGGCGGTGWGTRC-3**′**) targeting the V6–V8 regions. PCR amplicons were purified, index with barcodes and sequenced on a MiSeq Sequencing System (Ilumina) at the Australian Centre for Ecogenomics (ACE), The University of Queensland. Tailing samples from the glasshouse trial collected on days 155 and 282 were mixed, and microbial DNA extracted using the methods described above. The extracted DNA was then loaded for metagenomic sequencing at the ACE. Library preparation and quality control has been described previously^48^.

#### rRNA community analysis

Primer sequences were removed from forward de-multiplexed reads using cutadapt (ver. 3.2)^49^, with reads not containing primers discarded. Using QIIME2 (ver. 2022.2.0)^50^, reads were filtered, dereplicated and chimeras removed by DADA2 (--p-trunc-len-f 270)^51^. Taxonomy was assigned to the resulting amplicon sequence variants (ASV) by aligning each (classify-consensus-blast) against reference 16S rRNA sequences from GreenGenes2 (release 2024.09)^52^ and Eukaryote 18S sequences from the non-redundant SILVA database (release 138)^53^. ASVs that did not receive a classification at the phylum level or below, or that were classified as chloroplast or mitochondria, were discarded. ASVs with a relative abundance less than 0.01% in all samples were removed. To determine likely originating lineages, ASV sequences were aligned with mmseqs2 (ver. 15.6f452)^54^ against SSU rRNA sequences extracted from metagenome assembled genomes (MAGs) obtained from the present study, and from representative genomes from the genome taxonomy database (GTDB, R09-RS220)^55–57^. ASVs with an 98.7% identity match were considered species-level matches^58^. All statistical analyses were performed and visualizations generated in R (ver. 4.4.1).

To account for the compositional nature of the data, ASV counts were robust centered log-ratio (rclr)^59^ transformed with the decostand function in vegan (ver. 2.6-8)^60^ prior to principal-component analysis (PCA). PCA was performed using the rda function on species-level rclr-transformed^59^ counts with Euclidean distances. Beta-diversity Jaccard dissimilarity values (presence/absence) were also calculated using vegan. Environmental variables (pH, etc.) were fit to the ordinations using the envfit function from vegan.

For both the batch and glasshouse samples, abundances for species with an abundance ≥ 2% in at least one sample, or log_2_-fold difference of ≥ 5 according to edgeR (ver. 4.2.2)^61^ and an abundance ≥ 1%, were visualized as heatmaps. For glasshouse samples, log-fold values were calculated by comparing samples from a) the same amendment and replicate, e.g., OM + replicate 1, per day sampled, e.g., day 58 (and vice versa), and b) the same amendment, per day sampled (and vice versa); mean abundances were used when there were more than one sample in a group. Alpha diversity metrics Shannon (diversity) and Simpson (evenness), and number of observed taxa, were calculated using phyloseq (ver. 1.48.0)^62^. PCA figures were created with ggplot2 (ver. 3.5.1)^63^, ggnewscale (ver. 0.5.0) and vegan functions. Heatmaps were created using ComplexHeatmap (ver. 2.21.1)^64^. ASV metadata, sequences, taxonomy assignments and counts, and other associated data is provided in **Supplementary Table 4**.

### Metagenomic shotgun processing and analysis

#### Sample processing for metagenomics and metatranscriptomics

Metagenomic sequencing of the microbial DNA extracted from the glasshouse tailing samples was performed at the ACE. Library preparation and quality control has been described previously^48^. For metatranscriptomic sequencing, glasshouse tailing samples from days 155 and 282 were mixed and the corresponding composite inoculum sample used for RNA extraction. Microbial RNA was extracted by using a DNeasy PowerSoil Kit (Qiagen). RNA concentration was quantified using Qubit RNA BR. The TruSeq Total RNA Library Prep with Ribo-Zero Plus kit was used for RNA library preparation following the manufacture’s protocol. The library was sequenced on a NextSeq500 (Illumina, USA) platform at ACE (Brisbane, Australia).

#### Read quality-control and assembly

To identify and remove any potentially contaminating human DNA, raw reads were mapped to the human reference genome (GRCh38) with minimap2 (ver. 2.24)^65^, with those reads that mapped removed (less than 2% of the reads mapped; **Supplementary Table 5**). Low-quality reads were then identified and removed with Trimmomatic (ver. 0.39, ILLUMINACLIP:NexteraPE-PE:2:30:10)^66^. Quality controlled paired and singleton reads were assembled using metaSPAdes (ver. 3.15.4)^67^ with default parameters. Quality controlled reads were mapped onto their respective scaffolds with minimap2 as part of CoverM ‘make’ (ver. 0.6.1)^68^. Low-quality read mappings were removed with CoverM ‘filter’ (minimum identity 95% and minimum aligned length of 75%), and the number of remaining reads used to calculate the fraction of the DNA mapping to the assembled scaffolds. Finally, Nonpareil (ver. 3.4.1) was run on the quality-controlled reads using the k-mer alignment method to assess the fraction of the microbial community sampled by sequencing^69, 70^.

#### De-novo binning, quality evaluation and taxonomy assignment

The assembled sample was binned using the metagenomic binning pipeline Aviary (ver. 0.6.0, unpublished, R. Newell, https://github.com/rhysnewell/aviary). Briefly, Aviary first maps reads to the assembly with minimap2 (ver. 2.17) as part of CoverM (ver. 0.6.1) to obtain differential coverage information for each assembly. Using this coverage information, metagenome contigs were then binned using the Maxbin (ver. 2.2.7)^71^, MetaBAT (ver. 0.32.5)^72^, MetaBAT2 (ver. 2.15)^73^, CONCOCT (ver. 1.1.0)^74^, Vamb (ver. 3.0.2)^75^, Semibin (ver. 1.0.3)^76^ and Rosella (ver. 0.4.2; unpublished, R. Newell, https://github.com/rhysnewell/rosella) binning methods with a minimum contig length of 1,500bp and minimum bin size of 200,000bp. An optimal, non-redundant set of bins produced from the various binning tools were selected by DAS Tool (ver. 1.1.2)^77^. The completeness and contamination of all, 254 non-redundant bins were calculated by CheckM (ver. 1.1.3)^78^ and CheckM2 (ver. 1.0.1)^79^ (**Supplementary Table 6**), and taxonomy assigned to each using the classify workflow (‘classify_wf’) from the Genome Taxonomy Database Toolkit (GTDB-Tk; ver. 2.4.0; with reference to GTDB R09-RS220)^80, 81^. The non-redundant bins were then clustered and dereplicated using CoverM ‘cluster’ (precluster-method = dashing) with an ANI threshold of 97% and accounting for bin quality (--checkm-tab-table). Dereplication also yielded 254 bins, 71 of which of which had a quality **≥** 50 (calculated as the completeness – (3 X contamination) based on CheckM metrics); henceforth we refer to these 71 bins as the higher-quality metagenome assembled genomes (MAGs). The optimal growth conditions, e.g. pH and salinity, for the 71 MAGs were predicted using GenomeSPOT^27^.

To assess the novelty of the MAGs and for phylogenetic analysis, the closest reference genomes for each were identified from the class-level subtrees generated by GTDB-Tk during classification. Specifically, the closest two reference genomes by branch length in the respective subtree for each MAG were selected, along with one reference genome in the same order but not the same family (for maintaining the reference tree structure). The aligned set of bac120 marker proteins produced by GTDB-Tk for the 71 MAGs and corresponding reference genomes was used to construct a maximum-likelihood phylogenetic tree with IQ-Tree (ver. 2.2.3)^82^, with the best fitting model LG+F+R10 according to Bayesian Information Criterion, as chosen by ModelFinder^83^, and 1,000 ultrafast bootstraps.

#### Gene extraction and functional profiling

A non-redundant gene catalogue was constructed by first predicting protein-coding sequences (CDS) in the assembled scaffold using Pyrodigal (ver. 2.3.0)^84^, a Python library binding to Prodigal ^85^, in metagenomic mode. Sequences with start and stop codons, i.e., complete, were extracted using mfqe (ver. 0.5.0; B. Woodcroft, unpublished, https://github.com/wwood/mfqe). Complete protein sequences (438,202) were clustered at 100% protein identity using CD-HIT (ver. 4.8.1)^86^, with all members of each cluster required to have at least 80% of their sequence overlapping with the longest (seed) sequence. Dereplicated protein sequences (437,397) were functionally annotated using DRAM (ver. 1.4.6)^28^.

To estimate the gene abundance in the sample, quality-controlled reads of at least 140bp in length were aligned against the gene catalogue using DIAMOND blastx (--id 70, --query-cover 80, --min-score 40, --evalue 0.00001)^87^. The resulting read counts per gene were converted to reads per kilobase per million (RPKM) to account for gene length and metagenome size. Reads were then aligned to the set of 59 single-copy ribosomal marker genes packaged with SingleM (ver. 0.18.0)^88^ and RPKM values calculated. The RPKM values for genes extracted from the metagenomes were subsequently normalized by the mean of the 59 single-copy marker genes (in RPKM) to produce the final normalized RPKM value—an estimated percentage of the community with the gene, assuming one copy per genome.

Protein-coding sequences in all bins were also predicted using Pyrodigal in metagenomic mode. Protein sequences were functionally annotated and the completeness or presence/absence of functional pathways summarized with DRAM. In addition, organic and inorganic sulfur cycle proteins were annotated using HMSS2^30^. Because polysulfide, thiosulphate and sulfur (Psr/Phs/Sre) reductases belong to the functionally diverse Mo/W-bisPGD-binding (MopB) superfamily and are difficult to distinguish based on catalytic-subunit sequence similarity alone^31, 32^, candidate loci were further evaluated using a targeted classification approach (**Supplementary Note 2**). This approach integrated HMMER profile searches, phylogenetic placement, transmembrane topology prediction, signal peptide prediction, and gene-neighborhood analysis. Candidate catalytic subunits and associated B-like electron-transfer and C-like membrane-anchor proteins were classified by integrating local operon architecture with reference phylogenies spanning PsrA/PhsA/SrrA and related MopB-family enzymes. Finally, SSU rRNA sequences were identified and extracted with barrnap (ver. 0.9, https://github.com/tseemann/barrnap).

#### Generation of community profiles

The relative abundance of the dereplicated bins was calculated by first mapping the reads to each using CoverM ‘make’ and removing low-quality mappings with CoverM ‘filter’ (minimum identity 95% and minimum aligned percent of 75%). The mean coverage of each bin was then calculated with CoverM and the relative abundance of each, among those obtained, was calculated as its coverage divided by the total summed coverage of all bins. For visualization, relative abundances were scaled by the fraction of the (filtered) reads that mapped to the dereplicated bins.

The bacterial and archaeal community composition was also determined by classifying those raw reads corresponding to 59 single-copy genes using SingleM ‘pipe’, based on taxonomies derived from the Genome Taxonomy Database (GTDB) R09-RS220^55, 57^. SingleM ‘condense’ was used to produce a single OTU table containing the trimmed mean coverage across each lineage, calculated across all genes. SingleM ‘microbial_fraction’ was used to estimate the fraction of the reads that were microbial^89^. The relative abundance of each lineage was then calculated as its respective coverage divided by the total summed coverage.

#### Transcriptomics analysis

Metatranscriptomic reads were quality controlled with Trimmomatic (ver 0.39; ILLUMINACLIP:NexteraPE-PE:2:30:10, MINLEN:50). Ribosomal RNA was then detected and removed with RiboDetector (ver. 0.3.1)^90^. To identify the active microbial taxa, the remaining mRNA reads were mapped to the 254 dereplicated bins using CoverM ‘genome’ (ver. 0.7.0), with low-quality mappings filtered out (minimum identity 95% and minimum aligned percent of 75%). The number of reads mapping and transcripts per million (TPM) were then calculated with CoverM. To determine gene expression, reads were then mapped to the dereplicated catalogue of 437,397 gene sequences extracted from the assembled scaffold with HISAT2 (ver. 2.2.1)^91^, and the number of reads mapping and TPM calculated using CoverM (minimum identity 95% and minimum aligned percent of 75%).

## Data availability

The shotgun (metagenomic and metatranscriptomic) and the SSU rRNA amplicon read data for the samples described in this study, as well as the 71 higher-quality MAGs, have been deposited in the NCBI Sequence Read Archive (SRA) (Accessions: SRR33329622, SRR31309736–SRR31309750, SRR31311414) and Genbank, respectively, under the bioproject accession PRJNA1184692.

## Acknowledgements

The work is financially supported by the Australian Research Council Linkage project funding (LP190100975), Rio Tinto Ltd. And Queensland Alumina Ltd.

J.Z. acknowledges the support of Australian Research Council DECRA Fellowship (DE240100822).

