## Supplementary material for "A self-amplifying microbial–abiotic sulfur relay drives field feasible bauxite residue remediation"

**Supplementary Notes**

**Note S1**

Method (Modified Sobek ANC test ^1^)

1) Weighing 1g dried and finely grinded (prefer <75μm) in 50ml centrifugation tube, add 25ml 1M HCl (with 2 blanks), record the weight in the table in section 3.
2) Cover the tubes with white cap (stored in the hot block cabinet below the sink). The covered tube is heated to 90 ℃ for 2 h in heating block to digest and cooled at room temperature for 1h.
3) 10mL DI water is used to flush any sample adhering the tube wall to the bottom.
4) The sample was centrifugated (4000rpm, 10min), and the supernatant was collected and filtered (filter paper No 41 or 3).
5) 30ml supernatant was transferred to the titration tube and back titration using 2M NaOH to pH 7.0 and 8.3.

**Note S2** **Targeted** **MopB analysis**

This describes the targeted analysis used to identify and classify putative polysulfide (Psr), thiosulfate (Phs) and related sulfur reductase (Srr) loci from the metagenomic genome bins. The workflow was designed to avoid assigning canonical PsrABC function from catalytic-subunit hits alone, because PsrA, PhsA, SrrA and several other Mo/W-bisPGD catalytic subunits belong to the broader MopB/DMSO reductase superfamily and can be difficult to distinguish from one another using sequence similarity alone. Candidate loci were therefore interpreted using a combined framework incorporating profile-HMM discovery, genomic neighbourhood structure, membrane-subunit topology, TAT signal prediction and phylogenetic placement against curated reference sequences. The code developed for this analysis is available at <https://github.com/julianzaugg/PsrABC_prediction> (commit 9f4d635).

### Candidate discovery using dual profile-HMM gates

Protein-coding sequences in all bins were first predicted using Pyrodigal (ver. 2.3.0)^2^ in metagenomic mode. Candidate MopB catalytic subunits were then identified using two complementary profile-HMM discovery gates applied across all predicted proteins. The final candidate set was defined as the union of both gates, allowing broad recovery of Mo/W-bisPGD enzymes while retaining sensitivity for divergent Psr/Phs/Sre-like hits detected by HMSS2.

The first gate searched all predicted proteins against the Pfam-A Mo-bisPGD domain profile PF00384 using hmmsearch from HMMER 3 (ver. 3.4) with an E-value threshold of 1 × 10−5 ^3, 4^. PF00384 was used as a broad discovery profile because it detects the molybdopterin-binding domain shared by many Mo-bis(pyranopterin guanine dinucleotide) enzymes. It was therefore treated as a discovery gate rather than as evidence for PsrA function. The second gate searched all predicted proteins against the HMSS2^5^ PsrAPhsASreA.hmm profile at the same E-value threshold. This profile was used to increase sensitivity for Psr/Phs/Sre-like catalytic subunits that may be poorly recovered by the broader Pfam Mo-bisPGD profile. Candidates recovered only through the HMSS2 gate were retained for downstream analysis but flagged for manual review because the PsrAPhsASreA.hmm profile encompasses SreA-clade sequences that are treated as a distinct group in the reference-tree analysis.

### Genomic neighbourhood extraction and analysis

For each candidate catalytic subunit, the local genomic neighbourhood was extracted from the Pyrodigal GFF annotations. Up to 10 genes upstream and 10 genes downstream of each candidate were retained. Neighbourhood protein sequences were written to a combined FASTA file and searched using HMMER against Pfam profiles selected to detect candidate electron-transfer and membrane-anchor subunits associated with Psr/Phs/Ttr-like reductase complexes. The following profiles were used with an E-value threshold of 1 × 10−5:

- PF12800 (NrfC-like), used to detect candidate PsrB-like iron–sulfur electron-transfer subunits;
- PF13247 (4Fe–4S dicluster), used as an additional broad profile for iron–sulfur electron-transfer proteins;
- PF03916 (NrfD family), used as a broad profile for NrfD-family membrane-anchor subunits, including PsrC-, PhsC- and TtrC-like proteins;
- PF14589 (NrfD_2), used as a higher-specificity profile for polysulfide reductase-like membrane subunits and treated as strong support for PsrC-like context; and
- PF00384, used to re-confirm the Mo-bisPGD domain in candidate catalytic subunits.

When multiple NrfD-family hits occurred in the neighbourhood of a candidate catalytic subunit, the representative membrane-subunit hit was selected by prioritising PF14589 hits over PF03916-only hits, then hits with resolved topology over unresolved topology, and then PsrC-like topology where present. Candidate loci were summarised according to local subunit composition, including ABC_complete, AC_only, AB_only and A_only architectures.

Additional HMSS2 profiles were searched against candidate and neighbourhood proteins for annotation and cross-validation. Catalytic subunit profiles included PsrAPhsASreA.hmm, SoeA.hmm and TtrA.hmm, whereas neighbourhood profiles included PsrBPhsBSreB.hmm, PsrCPhsCSreC.hmm, SoeB.hmm, SoeC.hmm, TtrB.hmm and TtrC.hmm. These HMSS2 results were retained as annotation fields in the final classification table, except for SoeA.hmm hits, which applied a score penalty in the absence of contradicting PsrC or PF14589 evidence.

### Transmembrane topology prediction and membrane-subunit interpretation

All NrfD-family neighbourhood proteins detected by PF03916 or PF14589 were analysed with DeepTMHMM (ver. 1.0) using default settings to predict transmembrane helix topology^6^. Membrane-anchor subunits were interpreted primarily from transmembrane helix count in combination with profile-HMM support and phylogenetic context. 8 predicted transmembrane helices were considered consistent with PsrC-like topology, based on the structurally characterised polysulfide reductase membrane anchor^7^. 9 predicted transmembrane helices were considered consistent with TtrC-like topology^8^, whereas five predicted transmembrane helices were considered consistent with PhsC-like topology^8, 9^. Proteins with 4–6 transmembrane helices were also reviewed for possible SoeC-like context, where supported by the catalytic-subunit phylogeny and HMSS2 annotations. Proteins with 10 or more predicted transmembrane helices were treated as potentially belonging to other reductase families.

### Signal peptide prediction

Candidate catalytic subunits were analysed with SignalP (ver. 6.0) using the “other” organism setting and slow-sequential prediction mode^10^. Predicted twin-arginine translocation (TAT/Tat-SPI) signal peptides were used as supporting evidence for periplasmic Psr/Phs/Ttr-like enzymes. Absence of a TAT signal was used to flag candidates that may represent cytoplasmic or non-Psr enzymes such as SoeA-like proteins. TATLIPO categorical predictions were treated as TAT-positive. Where TATLIPO calls were associated with missing or zero numeric TAT probabilities, the categorical call was retained but the numeric value was treated as unavailable for quantitative comparison.

### Reference sequence selection and phylogenetic analysis

Candidate catalytic subunits were placed into a reference phylogeny spanning PsrA/PhsA/SrrA and other major MopB/DMSO reductase families. Reference sequences were assembled from two sources. First, characterised representatives were retrieved from UniProt and NCBI for PsrA (P31075, Q72LA4), PhsA (P37600), TtrA (Q9Z4S6, WP_011715816.1), SoeA (D3RNN8), SreA (Q8NKK1), ArrA (Q7WTU0, Q5Y818), AioA (Q7SIF4), NapA (Q8EIJ1), TorA (P33225), DmsA (P18775), NarG (P09152), SerA (Q9S1H0), PcrA (Q47CW6) and FdhG/FdhH (P24183, P07658). Second, an expanded reference set was selected from the DMSORcompwoutMop dataset of Wells et al. (2023)^11^ (see <https://datadryad.org/dataset/doi:10.5061/dryad.18931zd29>), with approximately 25 representatives selected from each of the major DMSOR/MopB families including PsrA/PhsA/SrrA, TtrA/SrdA, bSreA/SoeA, ArrA/ArxA, NapA, NarG, DmsA, DorA/TorA and FdhG. Candidate and reference sequences were aligned using MAFFT (ver. 7.525) with --anysymbol --auto^12^. The --anysymbol option was retained to accommodate non-standard residues, including selenocysteine, present in some reference sequences. Poorly aligned columns were removed using trimAl (ver. 1.5.1) with -gt 0.8 -st 0.001 -cons 60^13^.

Maximum-likelihood phylogenies were inferred with IQ-TREE (ver. 3.1.1) using ModelFinder for automatic model selection^14-16^. Branch support was assessed using ultrafast bootstrap approximation and SH-aLRT tests (-bb 1000 -alrt 1000). Candidate clade assignment was performed using ETE 3 by identifying the nearest labelled reference sequence by patristic distance^17^. The nearest and second-nearest labelled reference clades, their patristic distances, the distance margin and distance ratio were retained for manual review. These diagnostics were used to flag uncertain placements but were not applied as hard classification thresholds.

### Evidence integration and classification

Candidate loci were classified using a tree-gated evidence-integration framework. In this framework, phylogenetic placement defined the primary enzyme-family group, whereas local genomic context and sequence features refined confidence only within compatible groups. This strategy was used because several non-Psr Mo-bisPGD enzymes can occur in A–B–C-like operons, and operon architecture alone is therefore insufficient to assign canonical PsrABC function.

Candidates lacking PF00384 support were classified as NOT_MoBisPGD_enzyme regardless of other evidence. PF00384-confirmed candidates were assigned to the following broad tree groups: (i) PsrA/PhsA/SrrA-compatible placements; (ii) ArrA/ArxA; (iii) bSreA/SoeA/aSreA; (iv) TtrA/SrdA; (v) known non-Psr DMSOR/MopB families, including ActB, AioA/IdrA, DmsA, DorA/TorA, FdhG, NapA, NarG, NasC/NarB and related families; and (vi) unresolved or unrecognised placements.

Assignments of TRUE_PsrA, LIKELY_PsrA and PsrA_or_PhsA required placement within the PsrA/PhsA/SrrA-compatible tree group. Within this group, confidence was refined using supporting evidence from TAT signal prediction, B-like iron–sulfur electron-transfer neighbours, NrfD-family membrane-subunit neighbours, PF14589 support and PsrC-like membrane topology. Candidates with compatible tree placement but insufficient subunit resolution were classified conservatively as PsrA_or_PhsA rather than as canonical PsrA.

Candidates placed in ArrA/ArxA, TtrA/SrdA or bSreA/SoeA/aSreA groups were classified as LIKELY_ArrA, LIKELY_TtrA_or_SrdA or LIKELY_SoeA_or_divergent, respectively. Candidates placed in known non-Psr MopB families were classified as LIKELY_nonPsr_MopB. If such candidates nevertheless retained A–B–C-like architecture, they were classified as LIKELY_nonPsr_MopB_operon_like and flagged for manual interpretation. This category captures loci in which operon architecture resembles PsrABC but the catalytic subunit is phylogenetically outside the PsrA/PhsA/SrrA-compatible group.

**Note S3 Description of genetic potential of the recovered 71 high-quality MAGs**

***Carbon metabolism pathway***

**1. Genetic potential of glycolysis pathway**

The presence of glycolytic pathways was determined by identifying metagenome bins containing one or more of seven known glycolysis pathways. These include:

1. Entner-Doudoroff pathway: Converts glucose-6-phosphate (G6P) to glyceraldehyde-3-phosphate (G3P) and pyruvate.
2. Embden-Meyerhof pathway: Converts glucose to pyruvate or G6P to pyruvate.
3. Core glycolysis module with the non-oxidative Pentose Phosphate pathway: Converts glucose to pyruvate.
4. Core glycolysis module with the Pentose Phosphate pathway: Converts G6P to pyruvate.
5. Core glycolysis module with the non-oxidative Pentose Phosphate pathway: Converts fructose-6-phosphate (F6P) to pyruvate.
6. Pentose Phosphate pathway integrated with glycolysis: Converts G6P to G3P, followed by the glycolysis core module converting G3P to phosphoenolpyruvate.
7. C4-dicarboxylic acid cycle with NADP-malic enzyme, integrated with the TCA cycle.

**2. Genetic potential of heterotrophic growth**

Heterotrophic potential was inferred based on gene content in metagenome-assembled genomes (MAGs). This included the presence of genes associated with central carbon metabolism such as glycolysis, at least one pathway for carbon substrate utilization (e.g. gluconeogenesis for organic acid utilization or *acs*/*pta-ackA* for acetate utilization), and the tricarboxylic acid (TCA) cycle with completeness > 50%, as well as genes for the degradation of sugars and polysaccharides^18-20^. Carbohydrate-Active Enzymes (CAZymes) were annotated using the CAZy database via DRAM (ver. 1.4.6)^21^. The identified CAZy families, including glycoside hydrolases (GHs), carbohydrate esterases (CEs), polysaccharide lyases (PLs), and auxiliary activities (AAs), are putatively associated with lignocellulose degradation^22, 23^. Marker genes associated with multiple carbohydrate utilization pathways were identified, suggesting the potential to process sugars such as fructose, xylose, and sucrose into intermediates that can enter central carbon metabolism. Glycolysis then converts these substrates into acetyl-CoA and subsequent energy metabolism.

Organic acid utilization was inferred from the presence of genes associated with gluconeogenesis, enabling the transformation of amino and organic acid substrates into intermediates such as pyruvate or oxaloacetate. In addition, genes associated with acetate metabolism including two key pathways: (1) acetyl-CoA synthetase (ACS), and (2) the acetate kinase (AckA)–phosphate acetyltransferase (Pta) pathway. These pathways can convert acetate to acetyl-CoA, which can then be oxidized in the TCA cycle, used for fatty acid synthesis, or converted to succinate through the glyoxylate cycle. Details regarding marker gene counts, pathway completeness, and heterotrophic status for each MAG are provided in **Supplementary Table 12**.

Analysis of carbon utilization potential across the 71 high-quality MAGs indicated that 55 MAGs encoded genes associated with the degradation of acetate, organic acids, sugars, or polysaccharides, with tricarboxylic acid (TCA) cycle completeness exceeding 50% in these genomes. Genes linked to acetate metabolism—including *acs* (acetyl-CoA synthetase) and the *pta–ackA* pathway—were detected in 51 MAGs, suggesting a widespread potential for acetate transformation. The potential for sugar utilization was inferred in 25 MAGs, all of which also encoded genes associated with acetate metabolism. Several of these genomes additionally contained genes linked to gluconeogenesis and polysaccharide degradation.

Genes putatively associated with polysaccharide degradation were identified in 15 MAGs (e.g., bins 3, 6, 8, 25, 55, 56, and 65), including glycoside hydrolase families linked to cellulose (GH5, GH16, GH20) and hemicellulose (GH5, GH16, GH30, GH43) breakdown. Among these, six MAGs (bins 9, 25, 27, 39, 41, and 65) exhibited the greatest inferred metabolic versatility, encoding pathways associated with the utilization of multiple carbon sources, including sugars, organic acids, and polysaccharides.

**3. Genetic potential of carbon fixation and mixotrophic growth**

The potential for carbon fixation and autotrophic growth was inferred across the 71 MAGs by evaluating the completeness of seven carbon fixation pathways^24^ using DRAM:

- 3-Hydroxypropionate bicycle
- Dicarboxylate-hydroxybutyrate cycle
- Hydroxypropionate-hydroxybutylate cycle
- Methanogenesis (CO₂ → methane)
- Reductive acetyl-CoA pathway (Wood–Ljungdahl pathway)
- Reductive citrate cycle (Arnon–Buchanan cycle)
- Reductive pentose phosphate cycle (Calvin cycle, CBB cycle)

We assessed both the pathway completeness (fraction of completeness) in combination with the presence or absence of key marker genes, as listed in **Supplementary Table 13**. MAGs containing the necessary marker genes all exhibited pathway completeness greater than 70%, indicating a strong genetic potential for autotrophic growth.

Five MAGs (bins 1, 14, 47, 54, and 63) encode key marker genes for the Calvin–Benson–Bassham (CBB) cycle, including: Phosphoribulokinase (PRK) [EC:2.7.1.19] and Ribulose-1,5-bisphosphate carboxylase/oxygenase (RuBisCO) [EC:4.1.1.39], The CBB cycle completeness in these bins exceeded 81%, suggesting an ability to fixate CO₂. Additionally, *Rhodobacteraceae* bin23 and *Geminicoccaceae* bin59 encode the complete 3-Hydroxypropionate bi-cycle (100%), further supporting their role in CO₂ assimilation. All seven MAGs (bins 1, 14, 23, 47, 54, 59, 63) also encode the genes for heterotrophic growth, including genes for acetate metabolism, e.g., *acs* (acetyl-CoA synthetase) or *pta-ack* (phosphate acetyltransferase-acetate kinase). *Rhodobacteraceae* bin23, *Limnoraphis* bin54, and *Geminicoccaceae* bin59 additionally encode genes for sugar degradation, while *Thioalkalivibrio* bin63 encodes genes associated with lignocellulose degradation. Together, these genomic features suggest a mixotrophic potential, enabling the use of both organic carbon and fixed CO₂. Detailed information is provided in **Supplementary Table 13**.

**4. Genetic potential of fermentation**

MAGs with putative fermentative potential were identified based on the presence of genes associated with glycolytic and fermentation pathways, including those linked to the production of alcohols, propionate, acetate, and lactate^25^. A total of 26 MAGs (bins 1–3, 8, 9, 14, 15, 19, 22, 25, 27, 33, 39, 46, 54, 56–60, 63, 65, 67–70) were inferred to have fermentative potential based on the presence of genes associated with glycolytic and fermentation pathways. Among these, 21 MAGs also encoded components of the electron transport chain, including oxygen-dependent terminal oxidases, suggesting the potential for facultative anaerobic metabolism (**Supplementary Table 14** and **Supplementary Table 15**). All 26 MAGs encoded genes associated with glycolysis, suggesting the potential to convert glucose or its intermediates (e.g., glucose-6-phosphate and fructose-6-phosphate) to pyruvate. Eight MAGs (bins 1, 2, 22, 39, 56–59) encoded genes consistent with acetate fermentation pathways. Limnoraphis bin54 additionally contained genes associated with alcohol fermentation, including alcohol dehydrogenase (EC: 1.1.1.1), which is involved in alcohol production. Furthermore, 21 MAGs (bins 3, 5, 8, 9, 14, 15, 19, 25, 27, 33, 39, 46, 54, 57, 60, 63, 65, 67–70) encoded genes associated with lactate fermentation pathways. Three MAGs (bins 12, 39, and 59) contained genes linked to propionate metabolism, suggesting the potential for its integration into central carbon metabolism via the TCA cycle. Detailed pathway annotations for the different fermentation types are provided in Supplementary Table 14.

***Genetic potential of respiration***

**1. Oxidative phosphorylation**

Bacterial respiration is broadly classified into aerobic and anaerobic respiration, with the latter further subdivided based on the specific electron acceptor utilized^26^. Variation in terminal oxidase composition was observed across the recovered MAGs, suggesting differences in adaptation to oxygen availability. *Glycocaulis* *spp.* bin22 and bin49, as well as *Glycocaulis profundi* bin55, encode complex IV: a high affinity cytochrome bd ubiquinol oxidase, which is a respiratory enzyme prevalent in various aerobic bacteria, particularly under low-oxygen conditions. Unlike other oxidases, cytochrome bd oxidase possesses a unique structure and function, facilitating electron transfer from ubiquinol to oxygen while generating water without producing reactive oxygen species (ROS). This enhances bacterial tolerance to oxidative stress, playing a crucial role in energy metabolism and survival under oxygen-limiting environments. In addition, 16 MAGs (bins 1, 3, 4, 7, 8, 23, 25, 28, 32, 47, 56, 59, 60, 63, 65, and 71) encoded cbb₃-type cytochrome c oxidases, another class of high-affinity terminal oxidases commonly linked to microaerophilic respiration.

In contrast, 42 MAGs (bins 1, 2, 4–6, 8, 9, 11, 12, 16–19, 23–28, 32–34, 37, 39, 41, 43, 46, 47, 49–51, 54–57, 59, 61, 62, 66, 67, 69, and 71) encoded low-affinity cytochrome c oxidases (prokaryotic type), suggesting the potential for aerobic respiration under relatively high oxygen conditions. Based on the distribution of terminal oxidases, MAGs were grouped according to inferred respiratory potential. MAGs encoding high-affinity oxidases (e.g., cytochrome bd or cbb₃-type oxidases) are likely adapted to low-oxygen environments, whereas those encoding only low-affinity oxidases are more consistent with aerobic respiration under higher oxygen availability. MAGs lacking Complex IV components may rely on anaerobic metabolic processes. The MAGs were grouped based on inferred respiratory potential derived from terminal oxidase gene content:

- Putative microaerophilic populations (encoding high-affinity oxidases, such as cytochrome bd ubiquinol oxidase or cbb₃-type cytochrome c oxidase): bins 3, 7, 22, 60, 63, and 65.
- MAGs with potential for aerobic and microaerobic respiration (encoding both low-affinity cytochrome c oxidases and high-affinity oxidases): bins 1, 4, 8, 23, 25, 28, 32, 47, 49, 55, 56, 59, and 71.
- MAGs consistent with aerobic respiration (encoding low-affinity cytochrome c oxidases only): bins 2, 5, 6, 9, 11, 12, 16–19, 24, 26, 27, 33, 34, 37, 39, 41, 43, 46, 50, 51, 54, 57, 61, 62, 66, 67, and 69.
- MAGs lacking Complex IV oxidases (suggesting potential reliance on anaerobic metabolic processes): bins 10, 14, 15, 21, 29, 30, 31, 36, 38, 40, 42, 44, 45, 48, 52, 53, 58, 64, 68, and 70.

**2. Phototrophic respiration**

A total of nine MAGs encoded components of the photosynthetic systems, with *Limnoraphis* bin54 containing genes for both photosystem I (PSI) and photosystem II (PSII). Studies reported that oxygenic phototrophs, including all cyanobacteria, utilize a combination of type I and type II photosystems where light drives oxidation of water, generating oxygen, protons, and reducing power^27^. The remaining eight MAGs (bins 23, 24, 26, 28, 29, 33, 46, 59) encode only PSII, predominantly found in phyla *Gemmatimonadota* and *Pseudomonadota*. A previous study suggested that *Gemmatimonas phototrophica*, an aerobic, anoxygenic phototrophic bacterium from the phylum *Gemmatimonadota* performs anoxygenic photosynthesis with PSII^28^. Bacteria that possess only PSII and lack PSI are relatively rare. This organism performs anoxygenic photosynthesis and relies on alternative electron donors with redox potentials more negative than water. These include reduced sulfur compounds (e.g., $H₂S$), molecular hydrogen ($H₂$), and organic substrates such as acetate^29^. Detailed annotations are provided in **Supplementary Table 15**.

**3. Nitrogen respiration**

The microbial community exhibited a broad genetic potential for nitrogen transformation (**Supplementary Table 16**), including pathways associated with dissimilatory nitrate reduction to ammonia (DNRA), denitrification, and nitrogen fixation. A total of 15 MAGs (bins 1, 7, 8, 15, 22, 23, 27, 34, 37, 46, 50, 56, 59, 62, 70) encode genes for anaerobic nitrate respiration (*napAB* or *narGHI*), suggesting the potential for the reduction of nitrate to nitrite. Among these, five MAGs (bins 7, 8, 15, 27, and 62) also encode genes associate with nitrite reductase (*nirBD* or *nrfAH*), consistent with the potential for DNRA. Genes associated with denitrification were detected in 24 MAGs, including key markers such as *nirS/K*, *norBC*, and *nosZ*, indicating a widespread potential for partial denitrification. In addition, nitrogen fixation genes (*nifHDK*) were identified in three MAGs (*Geoalkalibacter halelectricus* bin2, *Limnoraphis* bin54, and *Desulfuromonadales* bin58), suggesting the potential for diazotrophic activity.

**4. Sulfur respiration**

Putative genes involved in dissimilatory sulfur metabolism were identified using HMSS2^5^.

**Direct Cellular Contact via Outer Membrane Proteins (OMPs)**

Elemental sulfur ($S₈$) is predominantly present as hydrophobic, polymeric forms with limited bioavailability. Microbial utilization of S₈ has been proposed to involve direct cell–surface interactions, including thiol-mediated activation at the cell envelope^30^. Analysis of the genes from *Limnoraphis* bin54, and *Unclassified* bins 32, 60, and 63, showed that these bins possess the complete genetic repertoire for this mechanism, including components of the type I secretion system:

- ATP-binding cassette (ABC) transporter: *hlyB* (K11004)
- Membrane fusion protein (MFP): *hlyD* (K11003)
- Outer membrane protein (OMP): *tolC* (K12340)

These proteins form a continuous channel that facilitates the translocation of various substrates, ranging from small peptides to large S-layer proteins, in an unfolded state^31^. Studies have demonstrated that the *HlyD* and *TolC* components of the type I secretion system are upregulated during sulfur oxidation and reduction, suggesting their role in surface adhesion and the export of specialized OMPs involved in sulfur activation. This process is considered crucial for initiating sulfur oxidation and reduction pathways^32^.

**Polysulfide as the electron shuttle**

Under neutral or alkaline conditions, the nucleophilic attack of S_8_ by HS^−^ results in the nucleophilic cleavage of S_8_ rings and the formation of small polysulfide molecules (Eq. S1)^30^. Polysulfide molecules can pass through the cell membrane via channels or other designated polysulfide-binding carrier proteins and react with the cytoplasmic sulfur transferases^30, 33^. Polysulfide remarkably enhances the bioavailability of $S₈$, and thus greatly accelerates the process rates for sulfur reduction or oxidation.

$S_{8}^{0}+ HS^{-}\rightleftharpoons S_{8} S^{2-}+ H^{+}$	(Eq. S1)

**Dissimilatory sulfur cycling organisms**

*Sulfide Oxidation to sulfane sulfur (S0) via SQR and FccAB in periplasm*

Genes associated with sulfide oxidation were identified, including those encoding membrane-bound sulfide:quinone oxidoreductase (SQR) and flavocytochrome c sulfide dehydrogenase (fccAB)^34, 35^. These enzymes are commonly associated with the oxidation of sulfide to intermediate sulfur species, including sulfane sulfur and elemental sulfur (S⁰), in the periplasm. Multiple SQR types are encoded across 21 MAGs:

SQR type I and III: encoded in bin1. Type I SQR is known for its dual role in detoxification and sulfide-dependent respiration.

SQR type II: encoded in bins 2, 4, 14, 16, 22, 33, 41, 46, 54, 55, 57, 59, 65, 67, and 71.

SQR type III: encoded in bins 12, 18, 19, and 45.

SQR type V: encoded in 61 MAGs, with transcriptional evidence supporting its expression in bins 4, 34, 47, 50, 54, 56, and 65. In addition to SQR, *fccAB* genes encoding flavocytochrome c sulfide dehydrogenase (FCSD) are encoded by *Unclassified* bin36 and *Longimicrobiales* bin65. However, no transcriptional evidence of *fccAB* expression was identified, suggesting its role may be limited under the environmental conditions sampled. SQR channels electrons from sulfide oxidation into the electron transport chain at the quinone level, conserving more energy, FCSD passes electrons to cytochrome c, which ultimately transfers them to oxygen, yielding lower energy conservation^36^. The broader distribution and expression of SQR compared to FCSD reflects the energy efficiency of SQR.

*Sulfane Sulfur Oxidation to Sulfite via SDO in the Periplasm*

Sulfur dioxygenase (SDO) catalyzes the oxidation of sulfanylglutathione (GSSH) into sulfite ($SO₃^{2}⁻$) and reduced glutathione (GSH). SDO plays a crucial role in sulfide detoxification and the oxidation of sulfane sulfur, which are vital processes in sulfur metabolism^37^. SDO is encoded in 67 out of the 71 MAGs, with exceptions in bins 5, 13, 20, and 35.

In genomes that also encode *fccAB* or SQR, the recombinant system rapidly oxidizes sulfide to sulfane sulfur. This sulfane sulfur spontaneously reacts with reduced glutathione (GSH) to produce glutathione persulfide (GSSH), which is subsequently oxidized by SDO to generate sulfite. Alternatively, studies suggest that in some *Acidithiobacillus* species, extracellular sulfur is activated by membrane proteins, converted into persulfide, and further oxidized by SDO into sulfite and thiosulfate^38^.

*Sulfane sulfur oxidation to sulfite via hdr and dsr in cytoplasm*

Genes associated with two key pathways for sulfur oxidation to sulfite in cytoplasm were found: the Reverse Dissimilatory Sulfite Reductase (rDsr) system and the Sulfur Oxidation (sHdr) system^39, 40^. The rDsr system is found in *Rhodocyclaceae* bin1 and *Thioalkalivibrio_A sulfidiphilus* bin47, both of which encode essential components, including *oxDsrAB*, *oxDsrC*, *oxDsrMKJOP*, *dsrEFH*, and the iron-sulfur flavoprotein *dsrL*. These bins also contain post-translational modifiers, such as *dsrR* and *dsrS*, which are crucial for regulating Dsr enzymes, consistent with other sulfur-oxidizing bacteria^41, 42^.

Both *Thioalkalivibrio_A sulfidiphilus* bin47 and *Thioalkalivibrio* bin63 encode the complete sHdr complex (sHdrC1B1AHC2B2) and the lbpA1-dsrE-lbpA2 gene cluster^41, 43, 44^. Bin63 contains several auxiliary genes, including *sHdrI*, *sHdrJ*, *sHdrT*, and the transcriptional regulator *sHdrR*, which are rarely associated with the core sHdr complex, enhancing its sulfur oxidation potential^41, 45^. All three bins—*Rhodocyclaceae* bin1, *Thioalkalivibrio_A sulfidiphilus* bin47, and *Thioalkalivibrio* bin63—encode *tusA*, a key cytoplasmic carrier protein that transports sulfane sulfur and delivers it to sulfur oxidation pathways.

*Sulfite Oxidation to Sulfate in periplasm and cytoplasm*

Genes associated with three pathways potentially involved in sulfite oxidation to sulfate were identified. *oxAprBA*/*oxSat*, occurs in the cytoplasm, utilizing adenosine 5′-phosphosulfate (APS) as an intermediate^46^. This pathway is found in both *Thioalkalivibrio_A sulfidiphilus* bin47 and *Thioalkalivibrio* bin63. The other two direct pathways involve either sulfite oxidoreductase (*sorAB*) in the periplasm (encoded by bins 23, 57, 63, and 67)^47^ or sulfite oxidoreductase (*soeABC*) in the cytoplasm^48^ (encoded by bins 1, 23, 47, 63 and 71). However, to date, there is only genetic evidence supporting the activity of *soeABC*^49^.

*Thiosulfate Oxidation: soxABXYZ in periplasm*

The Sox system is a major pathway for thiosulfate, sulfide, sulfite, and elemental sulfur oxidation to sulfate in sulfur-oxidizing bacteria (SOB)^50^. Complete Sox systems, which catalyze thiosulfate oxidation to sulfate in the periplasm, consist of SoxYZ, SoxXA, SoxB, and SoxCD proteins^51, 52^. Additional Sox proteins, such as SoxV, SoxW, SoxO, SoxF, SoxE, SoxG, SoxH, SoxT1, SoxT2, SoxS, and SoxR, may also be present depending on the organism^5^. In the absence of the SoxCD sulfur dehydrogenase complex, thiosulfate oxidation proceeds via a truncated Sox pathway, producing sulfate and elemental sulfur. The elemental sulfur deposits can later be oxidized to sulfite through reverse dissimilatory sulfate reduction pathways (*rDsr* or *sHdr*). *Rhodocyclaceae* bin1 and *Thioalkalivibrio* bin63 encode the truncated Sox pathway (*soxAXYZB*), while *Rhodobacteraceae* bin23 encodes *soxYZ*, *soxB*, *soxCD*, *soxEF*, *soxG*, *soxH*, *soxT2*, and *soxV*, but lacks *soxAX*.

*Sulfur reduction*

Among the 71 high-quality MAGs, none encoded a complete canonical dissimilatory sulfate-reduction pathway, including reductive *sat*, *aprAB*, *dsrAB*, and *qmoABC*, or the *asrABC* sulfite-reduction system. Application of the targeted MopB classification framework (**Note S2**) identified two high-confidence loci encoding PsrA/PhsA/SrrA-clade catalytic subunits consistent with polysulfide or thiosulfate reductase function (**Supplementary Table 17**).

The highest-confidence locus was recovered from CSSed10-48 bin154 (family *Trueperaceae*), where a complete PsrABC-like architecture was confirmed by both B- and C-like subunit detection, with the membrane anchor carrying the high-specificity PF14589 domain and a PsrC-compatible eight-transmembrane-helix topology. A second locus in JARFUG01 bin58 (order *Desulfuromonadales*) also encodes a complete ABC-like architecture with a predicted TAT signal peptide; however, the absence of a PF14589-positive membrane subunit and weaker phylogenetic separation from alternative MopB clades reduce confidence in a canonical polysulfide reductase assignment for this locus. Both loci encode an adjacent rhodanese-type sulfurtransferase. In characterised polysulfide reductases such as that of *Wolinella succinogenes*^53^, the rhodanese domain facilitates polysulfide chain transfer to the molybdenum active site, and co-localised rhodanese-type enzymes have been implicated in polysulfide trafficking and sulfane sulfur homeostasis. The co-occurrence of this gene at both loci provides independent biochemical support for a polysulfide-related function consistent with their proposed role in the sulfur relay. The classification outcomes for all MopB candidates, including subunit architecture, TAT predictions, and phylogenetic placement diagnostics, are provided in **Supplementary Table 17**.

***Novel lineage of CPR***

The novel Candidate Phyla Radiation (CPR) lineage recovered from the metagenomes was restricted to a few distinct groups within the phylum *Patescibacteria*, including *Saccharimonadales* bin13, *Saccharimonadia* bin20, and *Paceibacteria* bin35. Like most CPR members, these MAGs are characterized by small genome sizes and significant gaps in core metabolic pathways, consistent with a symbiotic lifestyle^54, 55^. Studies have suggested that *Patescibacteria* adopt a symbiotic lifestyle, relying on other organisms for essential metabolites^56^. The degree of metabolic dependence varies among different groups within *Patescibacteria*, reflecting the diversity in their metabolic capabilities^56^.

All three CPR MAGs encode incomplete glycolysis and pentose phosphate pathways (PPP) in the conversion of glucose to pyruvate. Specifically, all three lack key glycolytic enzyme encoding genes like glucokinase (*glk*), which phosphorylates glucose to glucose-6-phosphate (G6P), and phosphofructokinase (PFK), which catalyzes the phosphorylation of fructose-6-phosphate (F6P) to fructose-1,6-bisphosphate (F1,6BP) using ATP. Despite these deficiencies, G6P can still enter glycolysis via a metabolic shunt, in which F6P is converted into glyceraldehyde-3-phosphate (G3P) through the non-oxidative PPP. However, all three MAGs have G3P to pyruvate deficiencies, with *Saccharimonadales* bin13 and *Saccharimonadia* bin20 lacking phosphoglycerate mutase (PGAM) and enolase, and *Rariloculaceae* bin35 lacking GAPDH. None of these MAGs encode the genes for the pyruvate and acetyl-CoA pathways. However, bin35 does encode *ldhA* and *dld*, which produce lactate D via fermentation pathways.

The recovered CPR MAGs are all high quality (CheckM2 completeness > 95%, contamination < 1%) yet display limited metabolic potential compared to previously reported CPRs. They lack genes for energy production, biosynthesis, central carbon metabolism, and carbon utilization. However, it remains possible that some of the missing functions are either present but highly divergent, or that these functions occur via unknown biochemical mechanisms.

**Supplementary Tables**

**Table S1**: Setup and monitoring of lab-scale incubation trials

| No | Group_name | Duplication | BR add (g) | $S_{8}$ (g) | Ground sugarcane mulch(g) |
| --- | --- | --- | --- | --- | --- |
| 1 | L_BR | 1 | 185 | 0 | 0 |
| 2 | L_BR | 2 | 185 | 0 | 0 |
| 3 | L_BR | 3 | 185 | 0 | 0 |
| 4 | L_3S | 1 | 185 | 5.55 | 0 |
| 5 | L_3S | 2 | 185 | 5.55 | 0 |
| 6 | L_3S | 3 | 185 | 5.55 | 0 |
| 7 | L_OM | 1 | 150 | 0 | 17.4 |
| 8 | L_OM | 2 | 150 | 0 | 17.4 |
| 9 | L_OM | 3 | 150 | 0 | 17.4 |
| 10 | L_OM3S | 1 | 150 | 4.5 | 17.4 |
| 11 | L_OM3S | 2 | 150 | 4.51 | 17.4 |
| 12 | L_OM3S | 3 | 150 | 4.51 | 17.4 |

| Group Name | Gove Pond 5 Bauxite Residue | | 40% v/v  Tree mulch (ml) | Mineral Amendment Blends | |
| --- | --- | --- | --- | --- | --- |
|  | Weight (g) | Volume (ml)* |  | $S_{8}$ (% w/w) | $S_{8}$ (g) |
| G_OM_1 | 700 | $490$ | $196$ | 0 | 0 |
| G_OM_2 | 700 | $490$ | $196$ | 0 | 0 |
| G_OM0.5S_1 | 700 | $490$ | $196$ | 0.5 | 3.5 |
| G_OM0.5S_2 | 700 | $490$ | $196$ | 0.5 | 3.5 |
| G_OM1S_1 | 700 | $490$ | $196$ | 1 | 7 |
| G_OM1S_2 | 700 | $490$ | $196$ | 1 | 7 |
| G_OM3S_1 | 700 | $490$ | $196$ | 3 | 21 |
| G_OM3S_2 | 700 | $490$ | $196$ | 3 | 21 |

**Table S2**: Setup and monitoring of glasshouse trial

|  | Substrates | |  |  |
| --- | --- | --- | --- | --- |
| Name | Octasulfur ($S_{8}$) | Cellulose | inoculum | Conditions |
| B_SCI | + | + | + | Anaerobic |
| B_SC | + | + | - | Anaerobic |
| B_SI | + | - | + | Anaerobic |
| B_S | + | - | - | Anaerobic |
| B_SCI | + | + | + | Open-air |
| B_SC | + | + | - | Open-air |
| B_SI | + | - | + | Open-air |
| B_S | + | - | - | Open-air |

**Table S3**: Batch experiments conditions

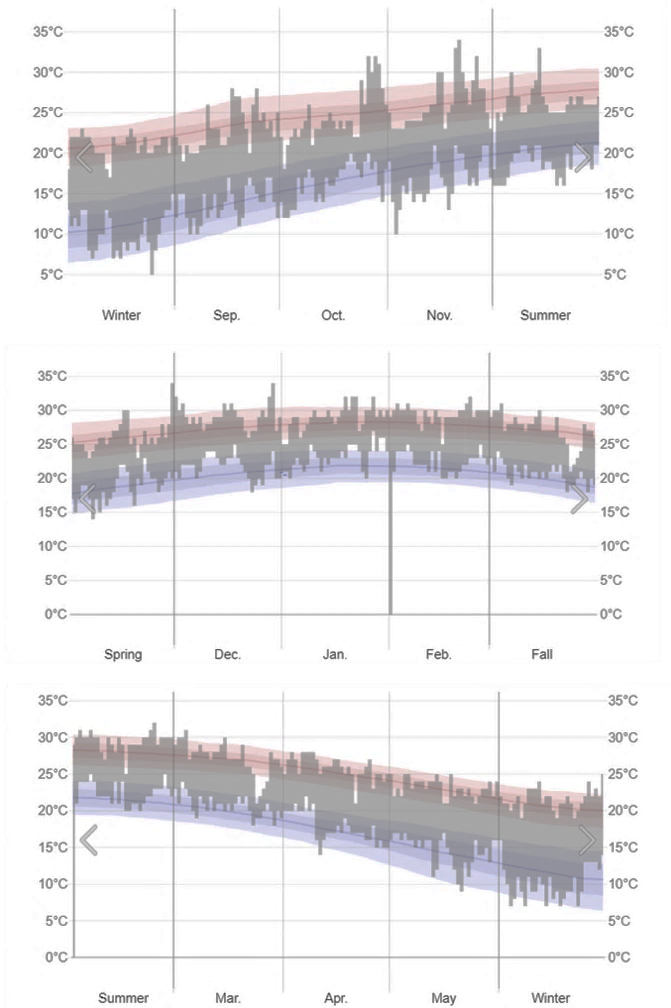
**Supplementary Figures**

**Figure S1**: Glasshouse experiment temperature started from Sep 2022 to May 2023 in Brisbane, Australia. Date from © WeatherSpark.com.

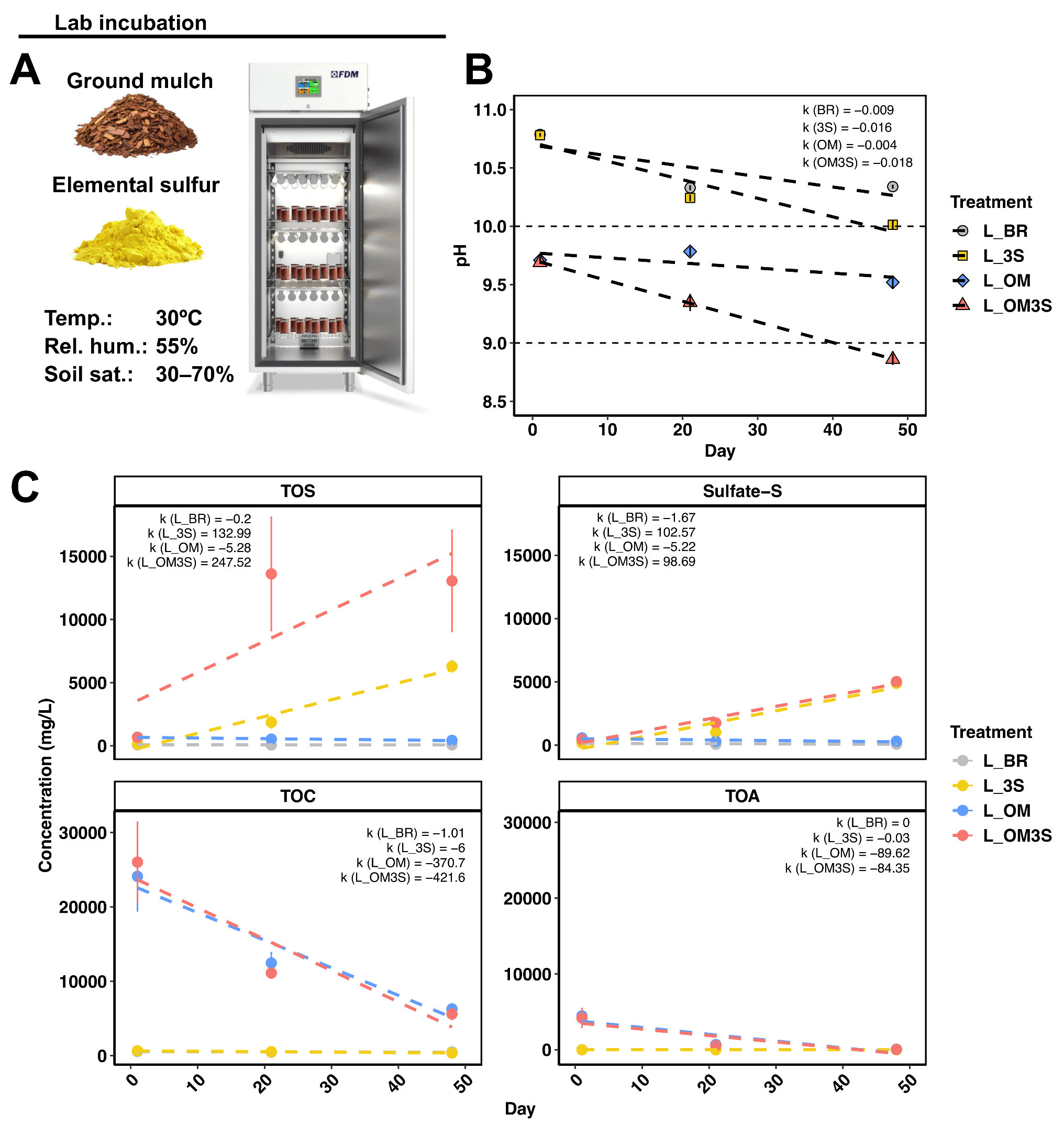

**Figure S2**: (A) shows the lab-scale experimental setup and conditions, featuring four treatment groups: (1) untreated bauxite residue (L_BR), (2) bauxite residue with 3% elemental sulfur (L_3S), (3) bauxite residue with 40% ground mulch as organic matter (L_OM), and (4) bauxite residue with both 40% ground mulch and 3% elemental sulfur (L_OM3S). (B) and (C) depict the dynamics of porewater geochemistry across the four groups over the 48-day incubation period, including total oxidative sulfur (TOS), total organic carbon (TOC), and total organic acid (TOA). Linear regression lines are fitted to each treatment group to show trends, with the slopes (𝑘) annotated on the plot to represent the rate of change in pH and the concentration of geochemical parameters over time for each treatment.

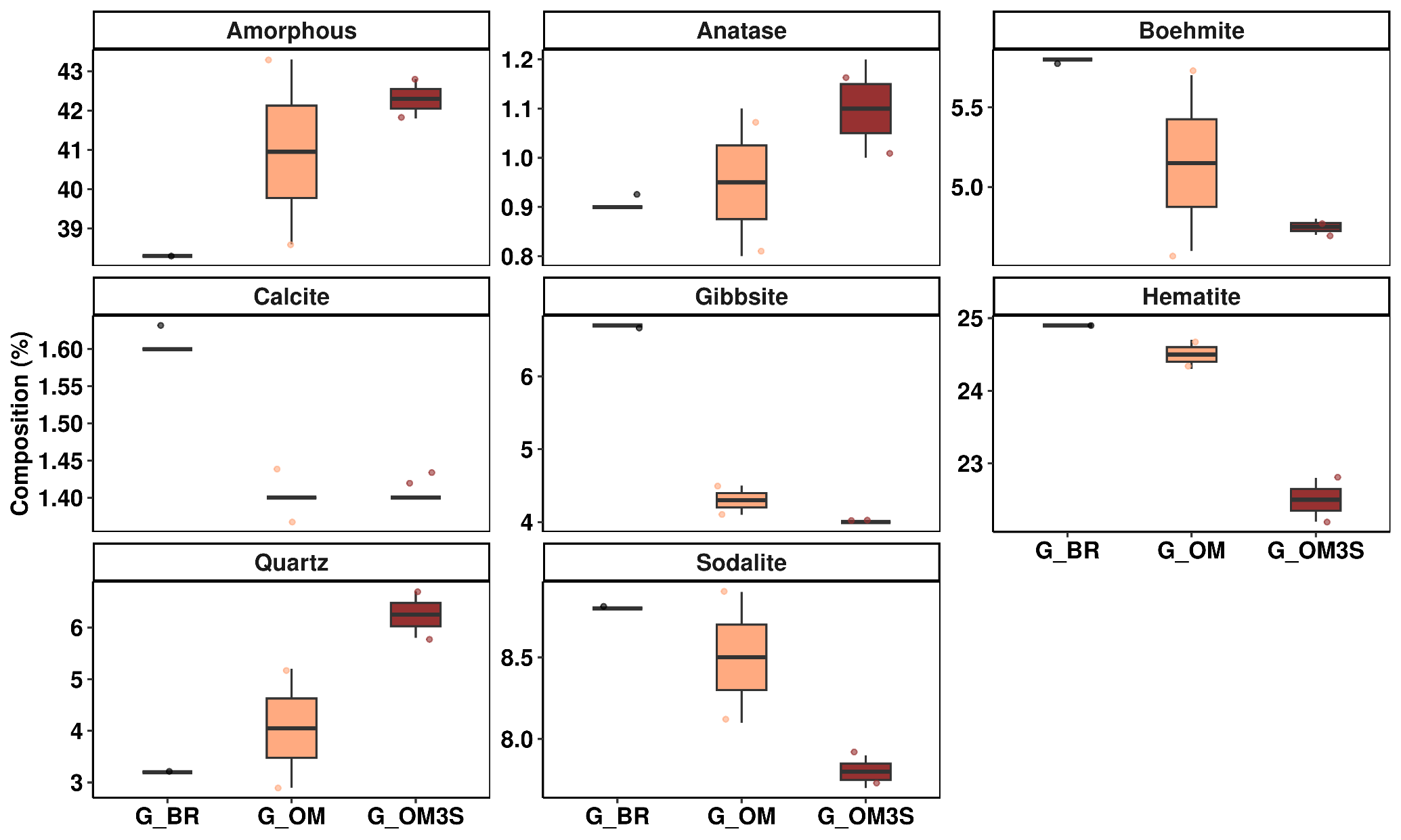
**Figure S3**: The mineral phase components from qXRD analysis in the Glasshouse trial. G_BR, bauxite residue without any amendments with inocula; G_OM, bauxite residue with 40% organic matter with inocula. G_OM3S, bauxite residue with 3% $S_{8}$ and 40% organic matter with inocula.

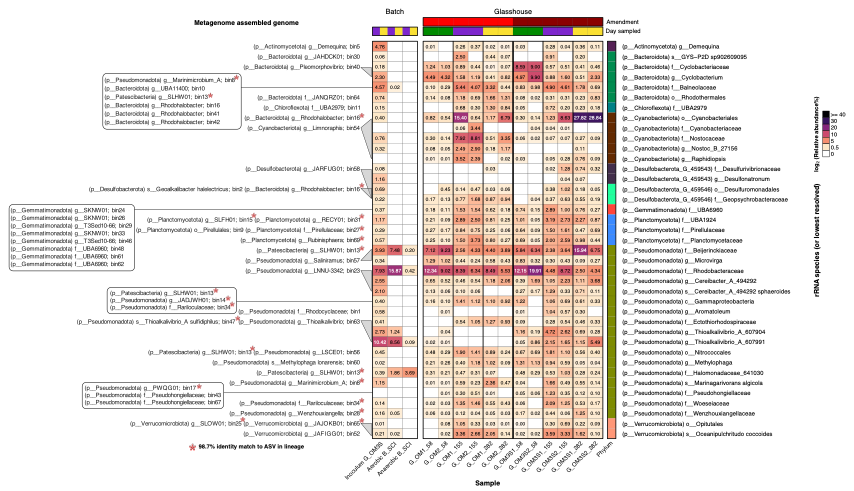

**Figure S4**: Heatmap showing the relative abundance of SSU rRNA amplicon sequence variants (ASVs) collapsed to the species level (or lowest resolved taxonomic level) across both composite G_OM3S inoculum samples collected from days 155 and 282, and replicate glasshouse trial G_OM and G_OM3S samples collected at days 58, 155, and 282. ASV sequences were aligned against SSU rRNA sequences extracted from the metagenome assembled genomes (MAGs) obtained from the present study to identify likely hosts, with a 98.7% identity considered a species-level match. An ASV sequence was included if it had either a species-level match to one of the MAGs, or its’ taxonomic classification matched that of the MAG (at least family level where possible). Likely ASV-MAG associations have been annotated to demonstrate consistency between the rRNA and metagenomic profiles. The color gradient represents log-transformed relative abundance, with darker hues indicating higher abundance levels. A suffix in a genus or species name, e.g., ‘_A’ in *Thioalkalivibrio_A*, is used in genome taxonomy database (GTDB) annotations to indicate polyphyletic groups, or to indicate the group has been subdivided based on taxonomic rank normalization according to the current GTDB reference tree. In the GreenGenes2 database, polyphyletic groups are indicated by a numeric suffix (e.g., ‘_607904’).

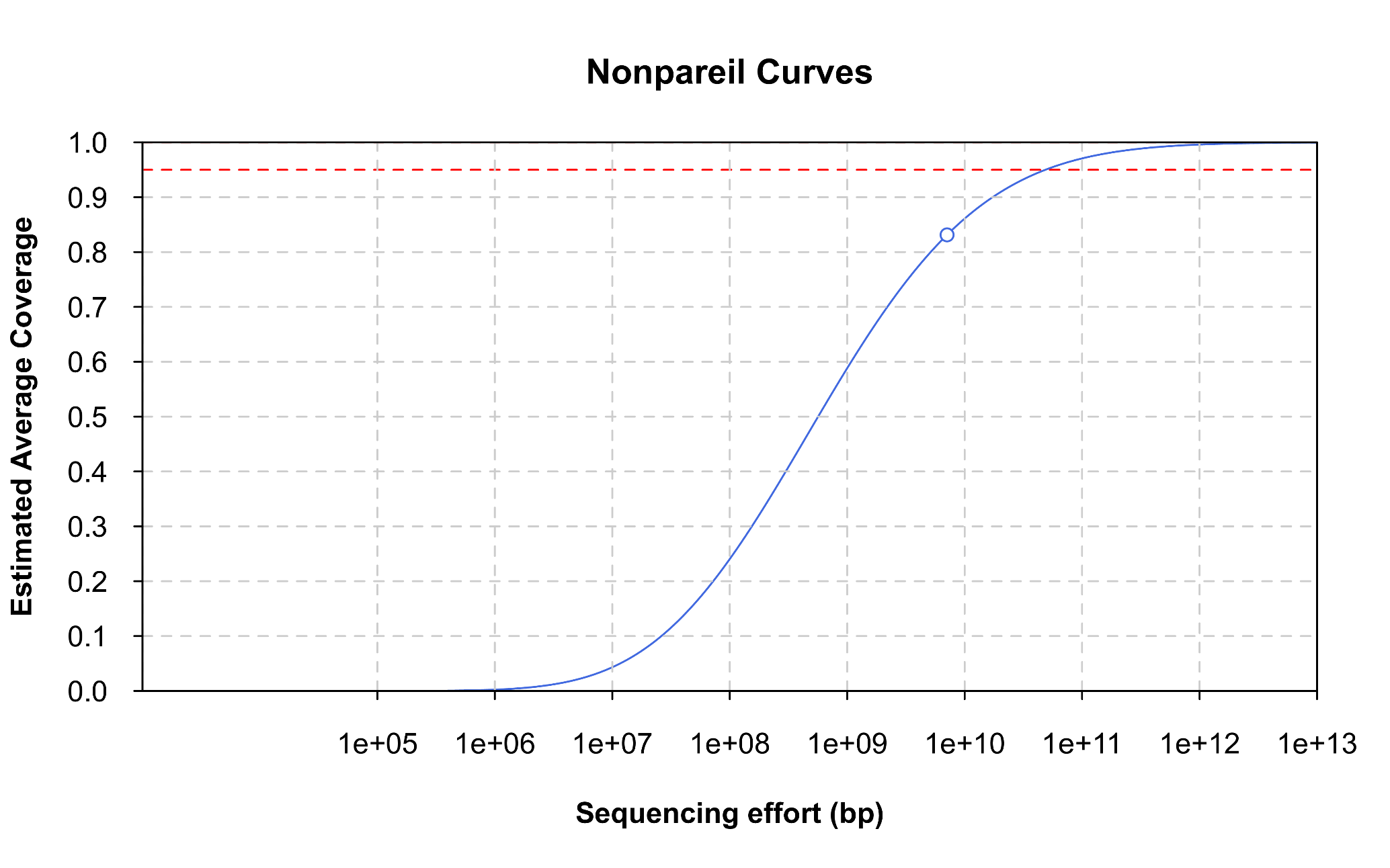

**Figure S5**: The estimated coverage (S-curve) and actual coverage (circles) for the shotgun metagenome sample evaluated in this study as reported by Nonpareil. The dotted red line represents 95% coverage.

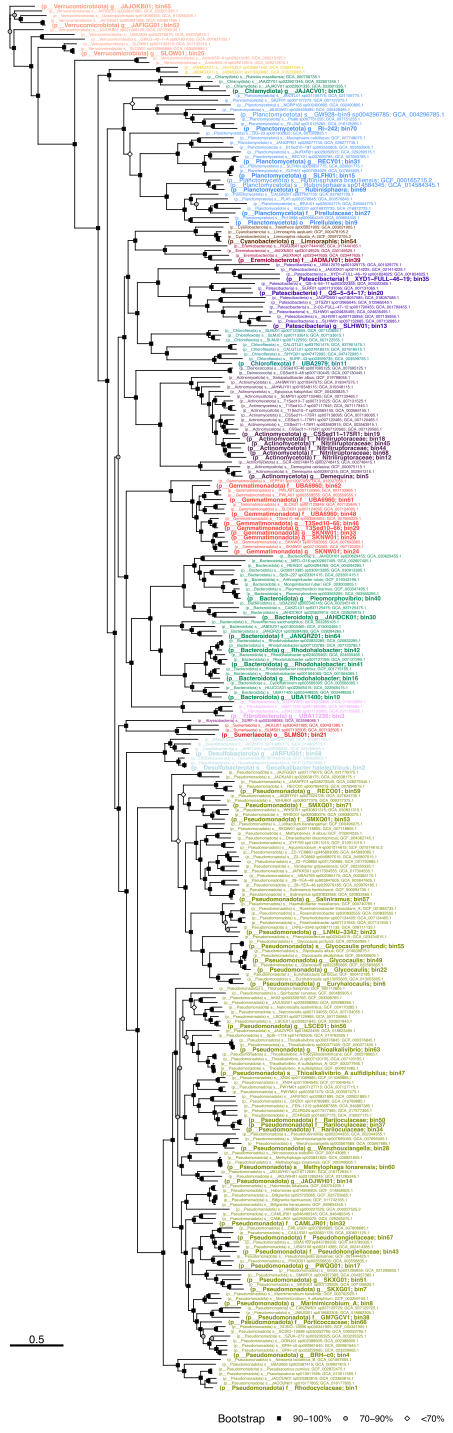

**Figure S6**: Maximum-likelihood phylogenetic tree based on aligned single-copy marker genes for 71 higher-quality metagenome assembled genomes (MAGs) assembled from the G_OM3S treatment (sampled at days 155 and 282), and 224 of their closest reference genomes. Leaf labels have been colored to indicate phylum, with MAG labels highlighted in larger bold text. Bootstrap values are represented by black squares for values of 90–100%, gray circles 70–90%, and white diamonds values <70%. The scale bar indicates the genetic distance.

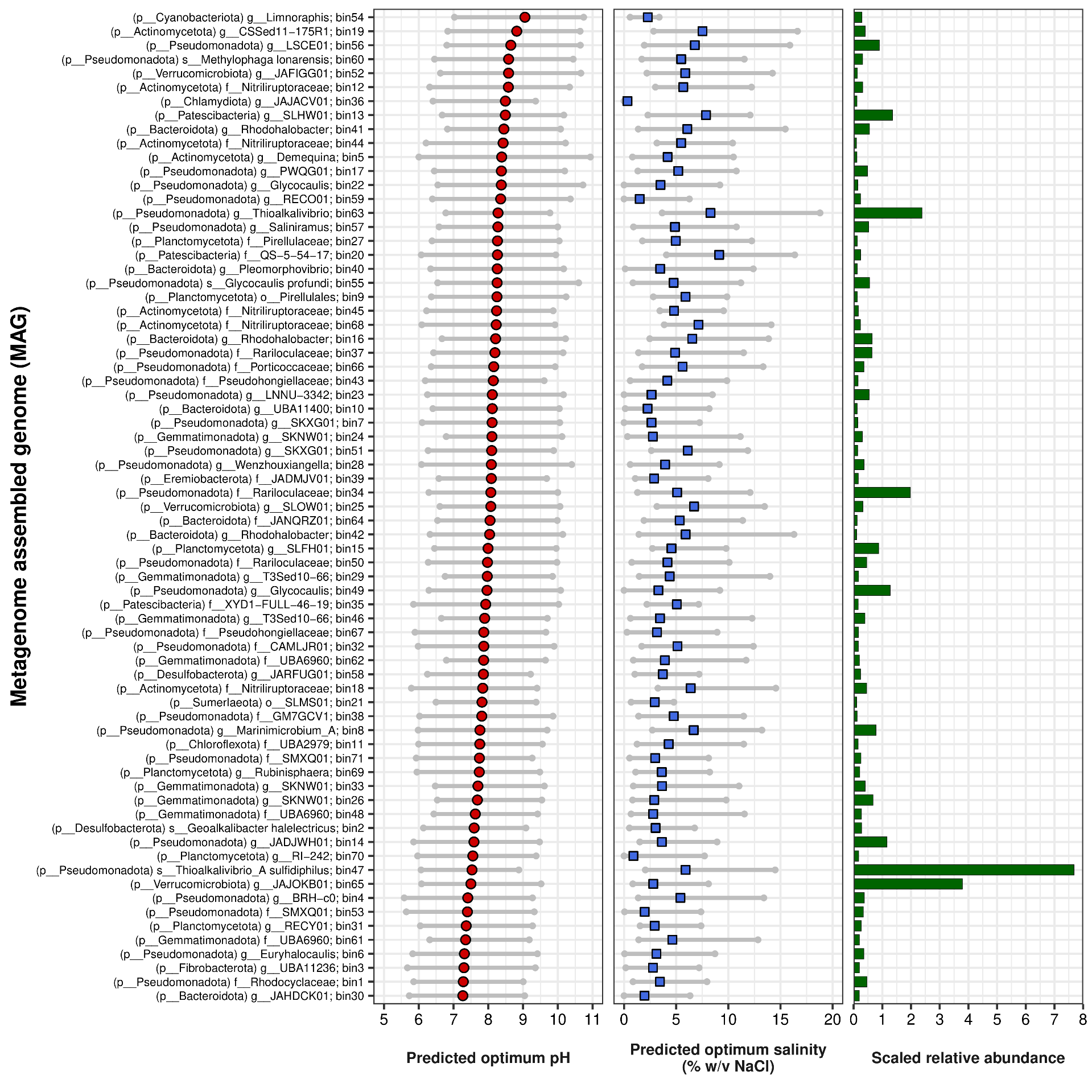

**Figure S7**: The optimum pH and salinity required for growth as predicted by genomeSPOT for the 71 higher quality MAGs. Scaled relative abundances are included for comparison—relative abundances scaled by the fraction of the (filtered) reads that mapped to the dereplicated bins.

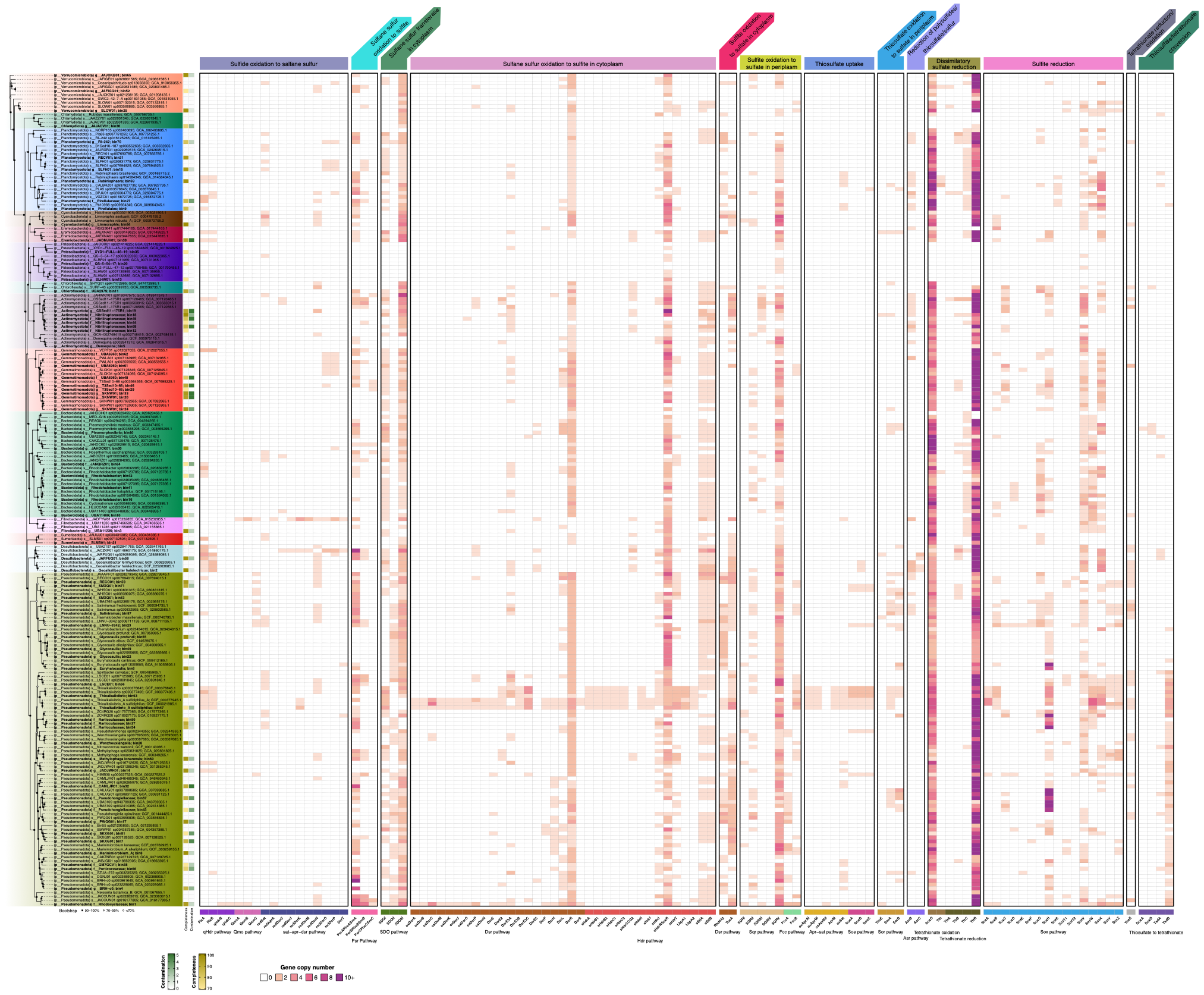

**Figure S8**: Heatmap illustrating the copy-number of sulfur metabolism-related genes (as identified by HMSS2) of the 71 higher-quality MAGs and closest reference genomes. The maximum-likelihood phylogenetic tree is shown on the left-hand side, with leaf labels colored by the phylum classification.

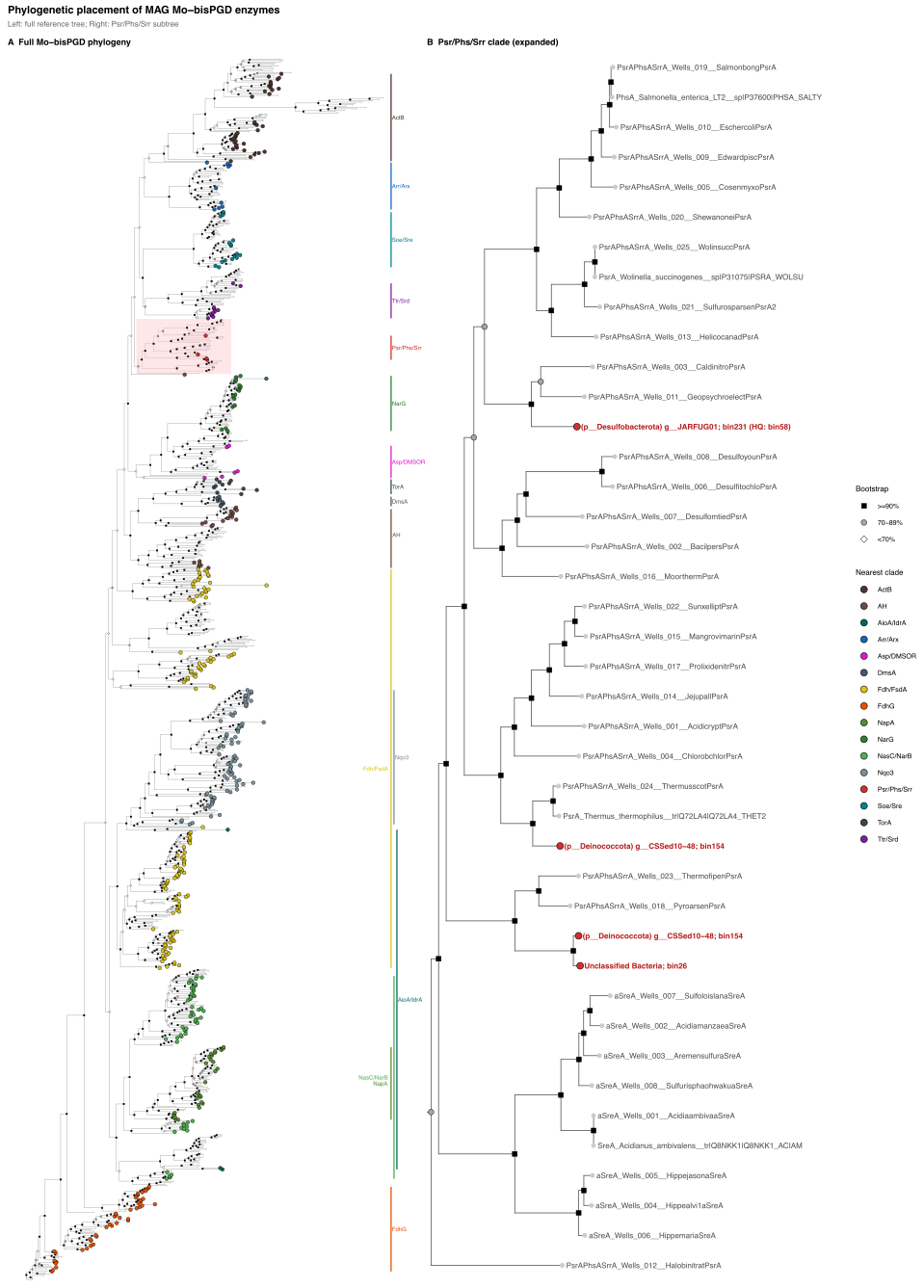

**Figure S9**: Maximum-likelihood phylogenetic tree of Mo/W-bisPGD-containing MopB-family catalytic subunits, inferred from genomic bins (including MAGs and lower-quality bins) and curated reference sequences. (**A**) Full phylogeny showing how bin-derived sequences (coloured points, coloured by nearest reference/clade) place relative to major MopB/DMSOR enzyme families; reference sequences are shown in grey. Coloured side bars mark major clade groupings among bin-derived sequences. The PsrA/PhsA/SrrA clade is highlighted here and expanded in panel B. (**B**) Expanded view of the PsrA/PhsA/SrrA clade, with bin-derived sequences labelled in red and reference sequences in grey. Phylogenetic placement was interpreted alongside operon structure, membrane topology predictions, TAT signal predictions, and neighbouring subunit analysis to assess evidence for PsrABC/PhsABC-like sulfur reductase architectures.

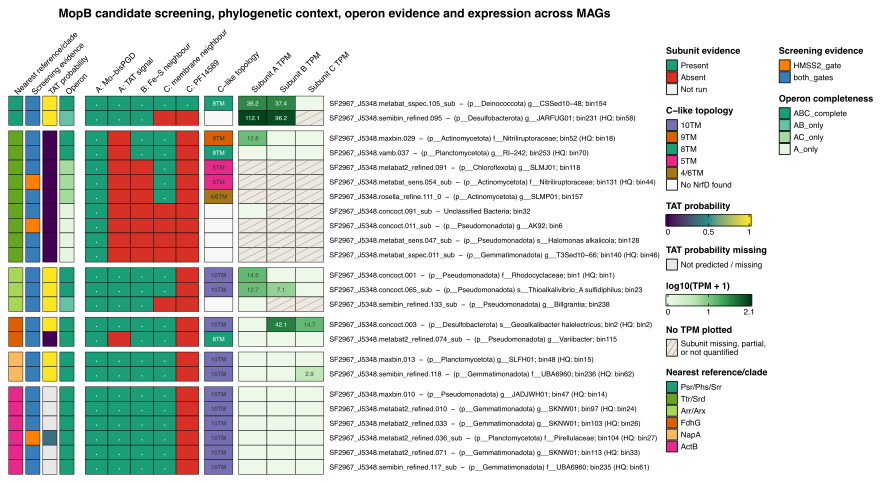

**Figure S10**: Heatmap summarising evidence supporting MopB-family catalytic subunits identified across genome bins. Each row represents a candidate locus selected from a bin based on combined phylogenetic and operon-context evidence. Annotations give the nearest reference/clade assignment (from phylogenetic placement), screening evidence (Mo-bisPGD domain [PF00384] and/or HMSS2 profile detection), TAT signal probability, and operon completeness. Presence/absence of supporting features is shown for the catalytic subunit (A: Mo-bisPGD domain and TAT signal), the neighboring Fe–S electron-transfer subunit (B), and the membrane-anchor subunit (C), including detection of the NrfD-family membrane domain PF14589. Predicted topology of subunit C is given as the number of transmembrane helices (TM). Expression levels for candidate subunits are shown as log10(TPM + 1) from metatranscriptomic read mapping. Hatched cells mark subunits that were missing, partial, or not quantified. Candidates placed outside the PsrA/PhsA/SrrA clade – including ArrA/ArxA-, TtrA/SrdA-, FdhG-, NapA- and ActB-related homologs – are shown for phylogenetic and operon-context comparison only and were not interpreted as evidence of canonical polysulfide or thiosulfate reductase activity.

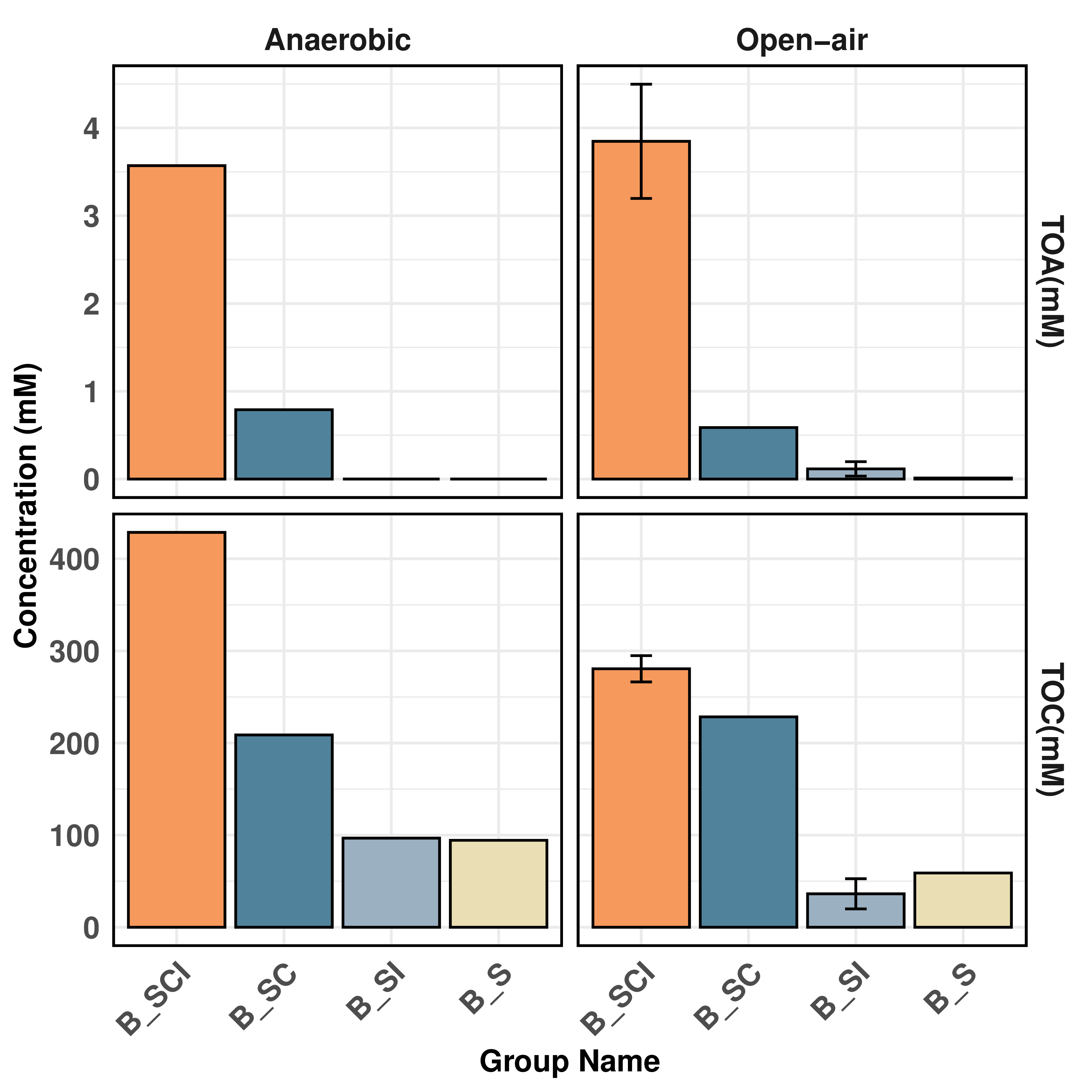
**Figure S11**: Batch experiments to uncover the dominant pathway for the S_8_ transformation and sulfuric acid production. Bar plot shows the total organic acid (TOA) and total organic carbon (TOC) in the final day of the batch after incubation. Batch study conducted in open-air flasks with oxygen flux and the second batch was conducted in anerobic conditions with serum bottle. The experiment group include different substrates as $S_{8}$ (B_S), $S_{8}$ + inoculum (B_SI), $S_{8}$ + cellulose (B_SC), $S_{8}$ + cellulose + inoculum (B_SCI).

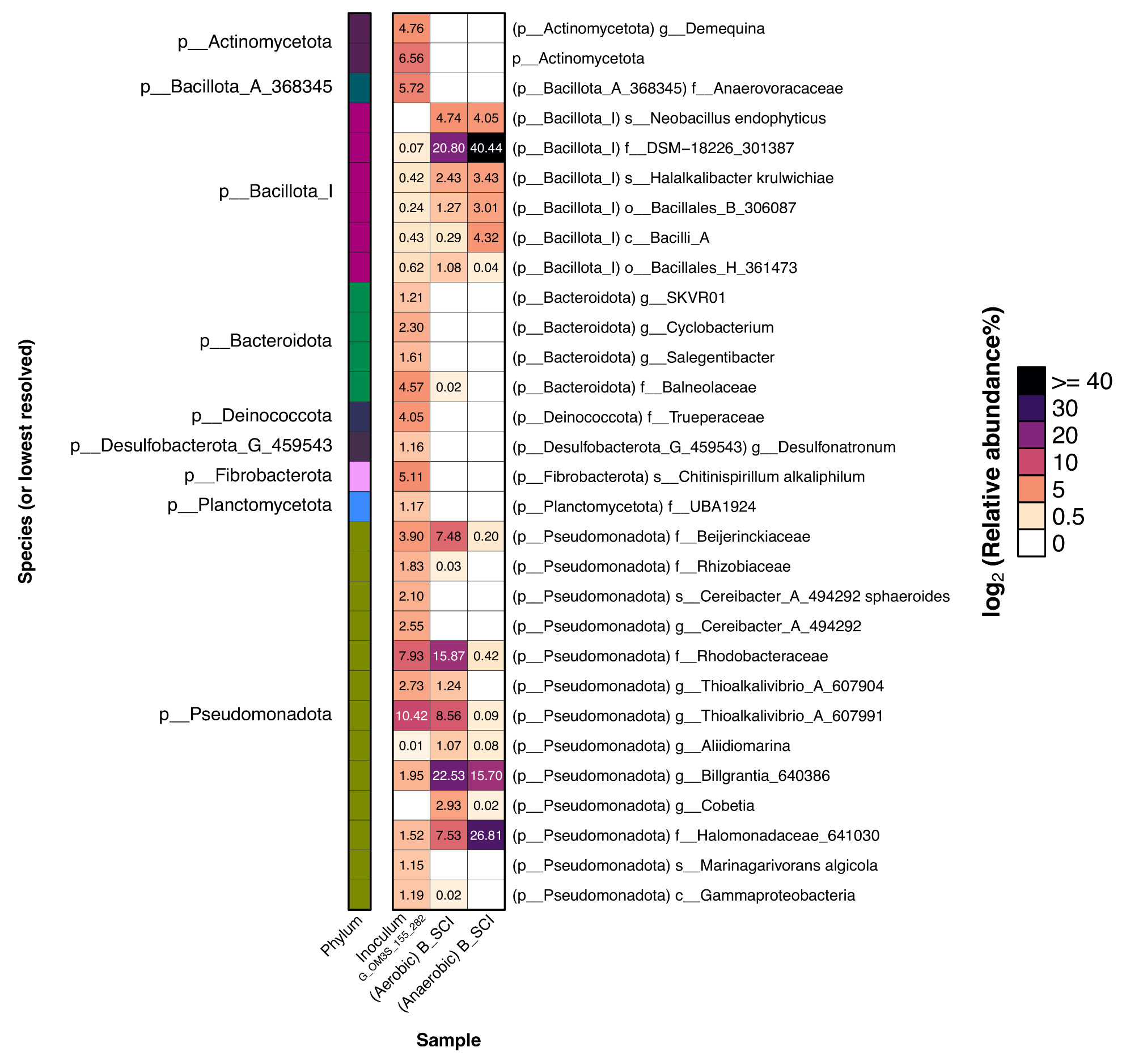

**Figure S12**: Heatmap of the rRNA relative abundances of microbial taxa from the batch incubation experiment samples: mixed G_OM3S samples from Days 155 and 282 as a microbial inoculum, and aerobic and anaerobic $S_{8}$ + cellulose + inoculum (B_SCI) samples.
